# Functional microRNA targeting without seed pairing

**DOI:** 10.1101/2025.03.03.641270

**Authors:** Matthew H Hall, Peter Y Wang, Thy M Pham, David P Bartel

## Abstract

MicroRNAs (miRNAs) associate with Argonaute (AGO) proteins to serve as guides, directing binding to partially complementary sites in mRNAs, ultimately causing post-transcriptional repression. Complementarity to the miRNA seed region (miRNA nucleotides 2–7) is typically both necessary and sufficient for repression. Here, we investigate unusual sites with extensive complementarity to the miRNA 3′ region (nucleotide 9 and onwards) but without complementarity to the seed. Top 3′-only sites bind as well as top canonical sites and impart similar repression, which can be further boosted by as few as 2–3 additional pairs to the miRNA seed. Despite these similarities, 3′-only sites have slower association and dissociation rates than seed-matched sites. They also impart different conformations to bound AGO–miRNA complexes than do seed-matched sites, and individual miRNAs differ substantially with respect to how well they bind their respective 3′-only sites. Thus, pairing to the seed is not required for binding and repression, or for a target to gain access to the 3′ region of the guide. Overall, for those miRNAs which recognize 3′-only sites, those sites are estimated to constitute 0.5–1% of the endogenous targetome, a proportion resembling that of other rare but functional site types, such as 3′-compensatory sites.

**GRAPHICAL ABSTRACT:** 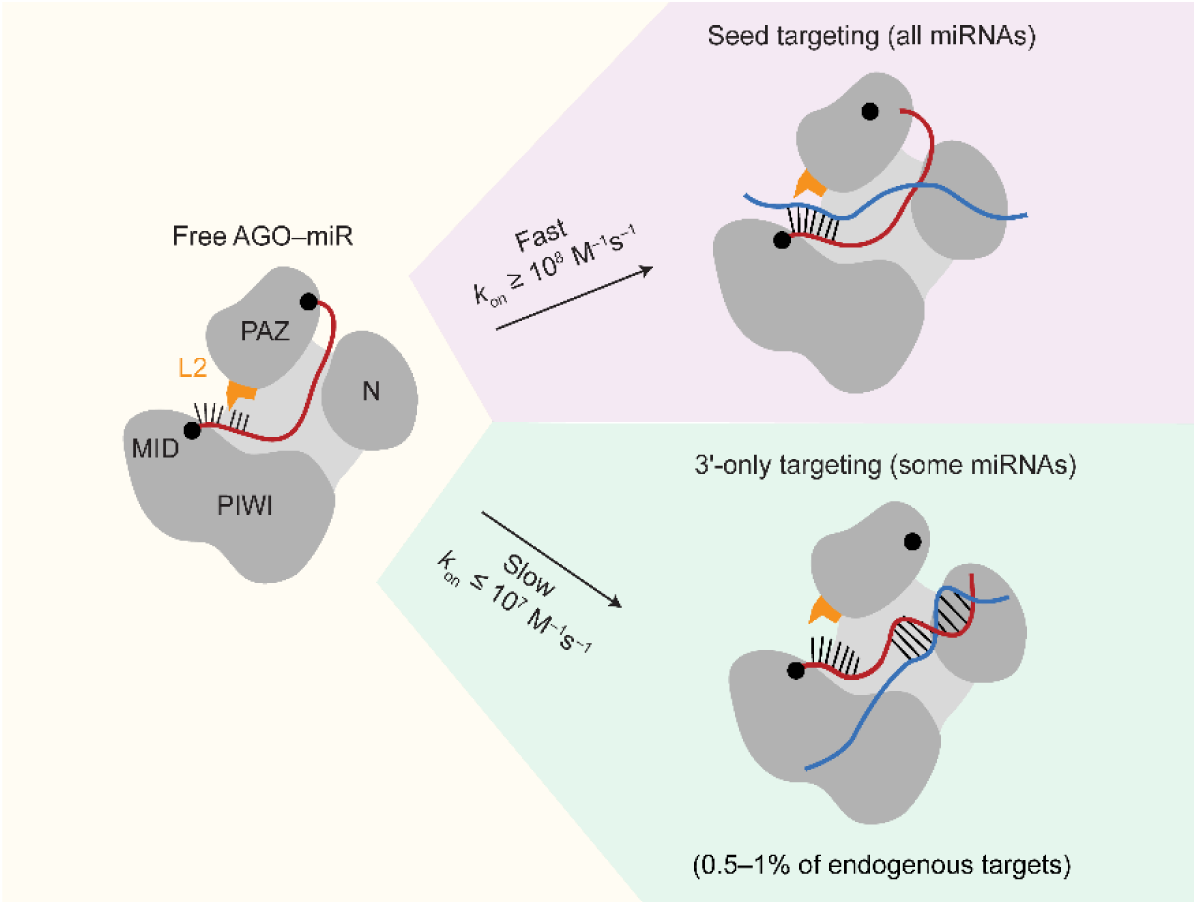

## INTRODUCTION

MicroRNAs are short (21- to 24-nt) regulatory RNAs that direct the post-transcriptional repression of target RNAs (1). After being processed from precursor hairpins, miRNAs associate with Argonaute (AGO) proteins to form AGO–miRNA complexes (AGO–miRs) (2). Within the AGO–miR, the miRNA acts as a guide RNA to bind partially complementary target sites, which in animal cells are most often in mRNA 3′ UTRs (1). AGO–miR bound to a target site recruits the adapter protein TNRC6, which in turn recruits the PAN2/3 and CCR4–NOT deadenylase complexes, which accelerate shortening of the poly(A) tail of the bound mRNA (3). Ccr4–NOT can also recruit inhibitors of translation initiation, which reduce translational efficiency of AGO– miR-bound mRNAs (3), although most posttranscriptional repression of the mRNA is due to accelerated deadenylation, which reduces translational efficiency in early embryos and destabilizes the mRNA in other contexts (4–10).

The target sites that are bound by AGO–miRs typically have perfect Watson–Crick complementarity to the miRNA seed (nucleotides 2–7), with either an additional pair to nucleotide 8 (creating what is called a 7mer-m8 site), or an adenosine (A) across from nucleotide 1 (creating a 7mer-A1 site), or both (creating an 8mer site) (1). Pairing to the seed is favored because the miRNA is spatially organized by contacts to AGO such that seed nucleotides 2–5 are accessible for pairing and pre-organized into a helical conformation that reduces the entropic penalty associated with duplex formation between miRNA and target in that region (11–17). Nonetheless, additional pairing to the miRNA 3’ region, which extends from position 11 to the end of the miRNA, can also play a role. For example, some pairing to the 3’ region can augment seed pairing to enhance targeting efficacy; such sites are called 3’-supplementary sites (18–23). Alternatively, particularly extensive pairing to the 3’ region can compensate for a mismatch or wobble in the seed pairing; such sites are called 3’-compensatory sites (18, 19, 21–27).

Comparative sequence analyses focusing on seed matches to broadly conserved miRNAs indicates that most mammalian mRNAs are evolutionarily conserved targets of miRNAs (26). Among the conserved seed-matched sites, only about 5% have signal for conserved 3’-supplementary pairing (19, 20, 26). The fraction of 3’-compensatory sites is even lower, estimated to comprise only 1–1.5% of conserved sites (20, 26). Experimental measurement of the binding affinities of different site architectures, and experimental measurement of changes in mRNA abundance and ribosome protected fragments after induction or perturbation of a miRNA both support and extend the results of these evolutionary analyses (4, 6, 8–10, 21–23, 27, 28). Canonical target sites, which feature perfect complementarity to the seed, constitute most consequential miRNA target sites, with site efficacy typically matching the expected hierarchy, with 8mer > 7mer-m8 > 7mer-A1 > 6mer (28). Across all types of target sites, the experimentally determined binding affinity between a target site and its cognate AGO–miR is highly predictive of the repression observed in cells (27, 28).

Argonaute RNA-bind-n-seq (AGO-RBNS) can simultaneously measure the relative AGO– miR binding affinities of all possible 8–12-nt RNA sequences (*k*-mers) (28). Such measurements have been acquired for six human miRNAs, which have revealed the relative binding affinities of many different target-site architectures of each miRNA. These affinity measurements can then be incorporated into a biochemical model that predicts mRNA repression as a function of total AGO–miR occupancy across all target sites present in the mRNA coding sequence and 3′ UTR (28). In an AGO-RBNS experiment, a library of RNA molecules is mixed with purified AGO–miR, allowed to reach binding equilibrium, and then bound molecules are isolated on a nitrocellulose filter that captures the AGO–miR and any associated RNAs (28). Bound RNA is extracted from the membrane, and libraries are prepared for high-throughput sequencing of both bound and input RNA samples. This procedure is repeated across a range of AGO–miR concentrations. For each AGO–miR concentration, the enrichment of a given site is calculated as the proportion of reads assigned to that site in the bound sample divided by the proportion of reads assigned to that site in the input sample. From the enrichment profile of a given site across the multiple AGO–miR concentrations, a relative *K*_d_ value is inferred that indicates how much better AGO– miR binds to the site compared to the no-site background (28).

For two of the six miRNAs analyzed by AGO-RBNS, namely miR-155 and miR-124, 10–11-nt *k*-mers matching the miRNA 3′ region were highly enriched in bound samples (28). Relative *K*_d_ values inferred from the enrichment profiles indicated that they bound their cognate AGO– miR with affinity comparable to that of 7-nt canonical sites. Binding to 10-nt 3′-only sites has also been detected in RBNS experiments with Drosophila Ago1 loaded with let-7, bantam, miR-184 and miR-11, with affinities depending on the miRNA and the register of 3′ complementarity, and the best sites binding about as well as 6mer canonical sites (29). However, for RBNS with either human or Drosophila AGO-miRs, whether these 10–11-nt *k*-mers were acting autonomously as 3′-only sites or whether they were only enriched because they were part of longer compound sites featuring both 3′ pairing and imperfect seed pairing could not be determined, because molecules with a given 10–11-nt match to the miRNA 3′ region but no contiguous downstream complementarity to the miRNA seed were too rare in the input library to calculate enrichments.

Other biochemical analyses report AGO–miR-21 and AGO–let-7 binding to targets with full 3’ complementarity but mismatches throughout the seed region. For the miR-21 seed-mismatched target, the affinity (*K*_d_ ∼ 1 nM;(23)) is >250-fold greater than background (*K*_d_ >250 nM;(22)); whereas for the let-7 seed mismatched target, the affinity is lower (22 nM; (22), consistent with the marginal but detectable AGO-RBNS enrichment of *k*-mers matching the 3′ region of let-7 (28). As for the highly enriched *k*-mers matching the 3′ regions of miR-155 and miR-124 (28), it is unclear whether short (e.g., 3-nt) matches to the seed in other registers contribute to this weak-to-moderate binding of let-7 and miR-21 sites that lack more extensive seed complementarity (22, 23). In cells, studies that aim to measure AGO–miR association with endogenous targets using cross-linking and sequencing report cross-linking to species with 3′ complementarity and little-to-no seed complementarity (30). However, because primary sequence context and pairing architectures strongly influence cross-linking efficiency, cross-linking cannot be used to accurately infer binding affinity or site efficacy of non-canonical sites (31).

We used modified AGO-RBNS, standard binding assays, and massively parallel reporter assays to determine binding affinities and cellular activities of 3′-only sites of miR-155 and miR-124 acting autonomously, which revealed 3′-only pairing configurations that bind with high affinity and confer repression in cells. Because functional 3′-only sites are substantially longer than seed-matched sites and do not occur for all miRNAs, they are relatively rare in the transcriptome. Despite the reduced scope compared to seed-based targeting, important 3′-only targeting interactions might ultimately be identified, especially as more miRNAs are identified for which this targeting mode is operative.

## RESULTS

### AGO–miR-155 and AGO–miR-124 bind 3′-only sites

To determine whether the *k*-mers identified in previous AGO-RBNS experiments corresponded to bona fide 3′-only sites, we conducted AGO-RBNS for miR-155 and miR-124 using a modified library design wherein half of the library molecules were programmed to contain an 8-nt match to the 3′ region of the corresponding miRNA (Fig 1A). The other half of the library molecules had a fully random-sequence region, which provided a diverse no-site background and a source of canonical and some noncanonical sites (Fig 1A). The presence of the programmed library molecules increased coverage of species with extensive 3′ complementarity, which allowed us to identify sufficient reads that featured stretches of 3′ complementarity not accompanied by any downstream seed complementarity and thereby determine relative *K*_d_ values for 3′-only sites acting autonomously, without the help of more than one nucleotide of contiguous pairing to the seed.

**Figure 1.**
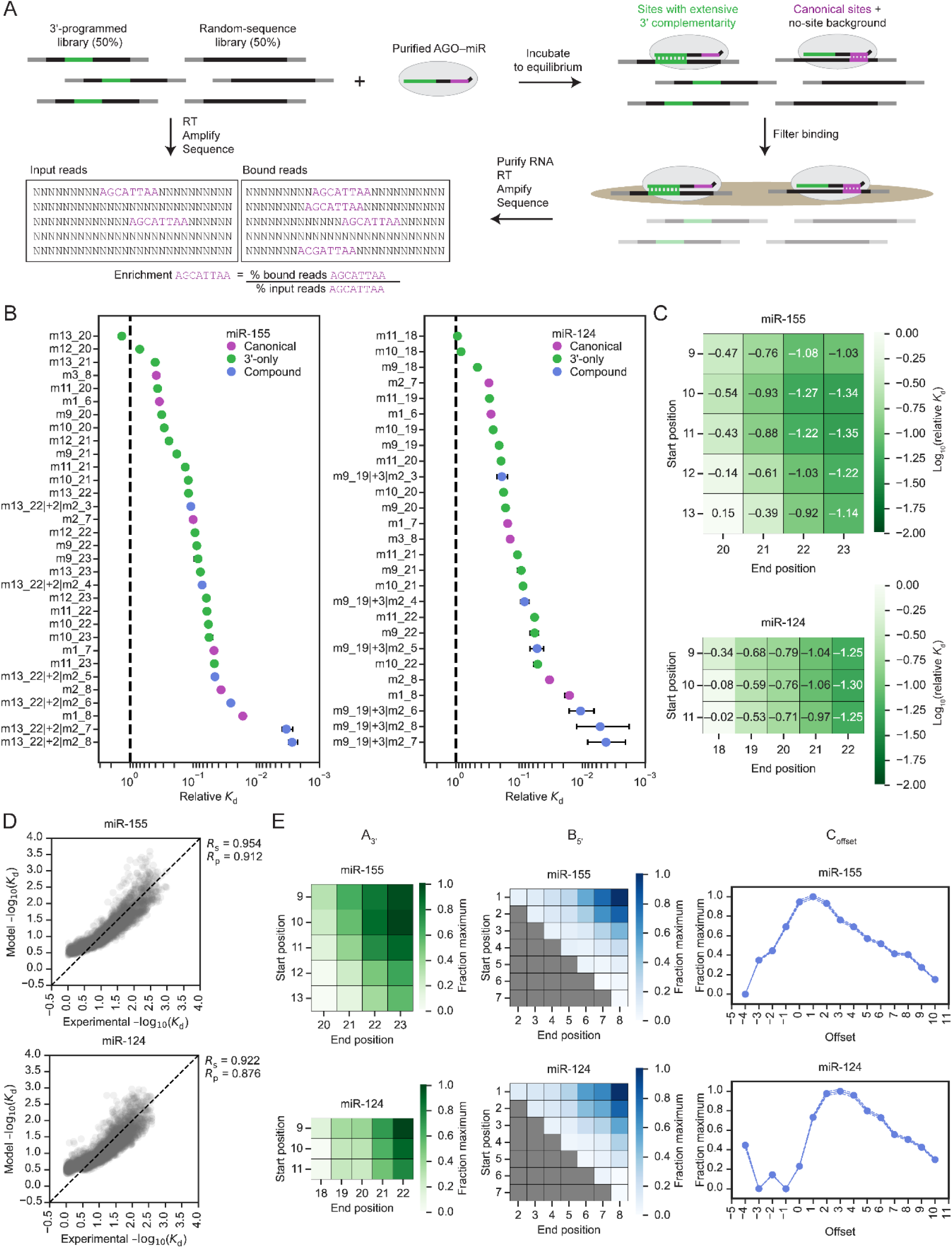
AGO-RBNS reveals high-affinity binding of AGO–miR-155 and AGO–miR-124 to 3′-only sites. (A) AGO-RBNS experimental workflow using a programed library to increase coverage of sites with extensive 3′ complementarity. The programed library had an 8-nt segment (green) that matched an 8-nt segment within the 3’ region of the miRNA (green). In the library, this segment was flanked by random-sequence positions (black), which were in turn flanked by primer-binding sites suitable for reverse transcription (RT) and amplification. See text for additional description of the RBNS procedure. (B) Relative *K*_d_ values inferred from AGO-RBNS enrichment profiles. Plotted are relative *K*_d_ values for canonical sites (purple), 3′-only sites (green), and select compound sites (blue) of miR-155 (left) and miR-124 (right). Error bars indicate 95% confidence intervals calculated as described (28). (C) Relative *K*_d_ values for different 3′-only site architectures. Heatmaps depict log_10_-transformed relative *K*_d_ values of different 3′-only sites of miR-155 (top) and miR-124 (bottom). (D) Relative affinities of compound sites of miR-155 and miR-124 are well fit by a multiplicative model in which negative log_10_-transformed relative *K*_d_ is a function of 3′ complementarity, seed complementarity, and offset. Shown are the model-predicted negative log_10_-transformed relative *K*_d_ plotted as a function of experimental negative log_10_-transformed relative *K*_d_ for sites of miR-155 (top) and miR-124 (bottom). Dashed black line shows *y* = *x*. Pearson (*R*_p_) and Spearman (*R*_s_) correlation coefficients for experimental versus model-predicted relative *K*_d_ are shown. (E) The influence of 3′-complementarity, seed complementarity, and offset coefficients on relative affinities of compound sites, as indicated by a multiplicative model for miR-155 (top) and miR-124 (bottom). At the left, heatmaps depict the relative magnitude of coefficients for different patterns of 3′ complementarity. In the middle, heatmaps depict the relative magnitude of coefficients for different patterns of seed complementarity. At the right, graphs plot the relative magnitude of coefficients for different offsets (top: miR-155, bottom: miR-124).

Each member of our programmed library for miR-155 had an 8-nt segment complementary to positions 13 through 20 of miR-155, as pairing to these positions of miR-155 appears to be more effective than pairing to other 8-nt segments of the miR-155 3’ region (28). Thus, the minimal 8-nt 3′-only site contributed by this library was complementary to nucleotides 13–20 of miR-155, designated as an m13_20 site. Likewise, our programmed library for miR-124 had a segment complementary to nucleotides 11–18, generating the minimal 3′-only site m11_18. For both miR-155 and miR-124, these 8-nt 3′-only sites did not bind detectably above background (Fig 1B). Nonetheless, when the 8-nt sites were extended by two or three additional nucleotides recruited from the random-sequence segments on either side of the programmed match, binding was detected. These 3′-only sites of 10–11 nucleotides bound about as well as 6-nt canonical sites for the respective miRNAs, and longer 3′-only sites with ≥12 nucleotides of perfect 3′ complementarity bound about as well as 7-nt canonical sites (Fig 1B). Longer 3′-only sites were generally stronger binders, and for 3′-only sites of equivalent length, the site with a more distal register was the stronger binder (Fig 1C). For example, m13_23 sites of miR-155 bound 4-fold better than m10_20 sites even though they were both 11-nt sites. This trend wherein sites of equivalent length but more distal register display stronger binding is also observed for other miRNAs in AGO-RBNS analyses using random-sequence libraries (28) and for the 10-nt 3′-only sites of fly Ago1–miRs analyzed by AGO-RBNS and conventional target-binding assays (29).

Despite their longer contiguous 3′ complementarity, sites that could pair to position 9 (e.g. m9_20) did not appear to bind better than otherwise identical sites that could not pair to position 9 (e.g. m10_20) (Fig 1C). For miR-124, sites beginning at position 9 (e.g. m9_20) were no better than equivalent sites beginning at position 10 (e.g. m10_20), and for miR-155, sites beginning at position 9 initially appeared to be worse binders than equivalent sites beginning at position 10. Others also observe that extending 3′ complementarity into the central region appears to be thermodynamically disfavored (23). We investigated this result by examining the flanking sequence preferences around 3′-only sites. If central pairing is indeed unfavorable, presumably due to steric hindrance because nucleotide 9 (and perhaps 10) of the miRNA are occluded by the central gate of AGO (32), then the thermodynamically most favored state for the guide–target duplex would be to leave nucleotide 9 unpaired, in which case, whether or not the nucleotide could potentially pair would be of no consequence. Therefore, an m9_20 site of miR-155 can instead be considered as an m10_20 site with a U residue flanking the m10_20 site (because nucleotide 9 of miR-155 is an A). When we re-ran our RBNS analysis pipeline focusing on the influence of the upstream flanking nucleotide on the relative affinity of m10_20 and other sites that started at position 10, miR-155 displayed a strong preference for A nucleotides immediately upstream of those 3′-only sites, whereas U and G nucleotides were particularly disfavored (Fig S1A). We conclude that 3′-only sites such as m9_20 of miR-155 appear to have worse affinity not because unfavorable central-region pairs actually form, but instead because to be classified as an m9_20 site, the nucleotide opposite position 9 must be a U, which is contextually disfavored, whereas the equivalent site with a contextually favored A opposite position 9 would be classified as an m10_20 site.

In sum, miR-155 and miR-124 bound 3′-only sites with at least 10 nucleotides of contiguous complementarity about as well as they bound 6-nt canonical sites, whereas longer and stronger 3′-only sites with more than 12 nucleotides of contiguous complementarity bound about as well as 7-nt canonical sites. This strong binding occurred even in the absence of complementarity to the miRNA seed, and longer and more distal 3′ pairing was associated with stronger binding, although position 9 remained unpaired.

### A match to the seed region as short as three contiguous nucleotides enhances affinity

Due to our increased coverage of sites with extensive 3′ complementarity, we were able to determine relative *K*_d_ values for many individual combinations of 3′ complementarity and seed complementarity. These different compound sites were classified based on three parameters: 3′ complementarity (e.g., m13_20), seed complementarity (e.g., m2_7) and the offset between the two (e.g., +2). Offset is calculated by comparing the number of nucleotides bridging the segments of 3′ complementarity and seed complementarity in the target site to the number of nucleotides bridging the corresponding segments of the miRNA (27). Positive offset values indicate that, compared to the miRNA, the target site has more nucleotides bridging the two complementary segments, whereas a negative offset value indicates that the miRNA has more nucleotides bridging the two complementary segments. For example, if a read featured m13_20 3′ complementarity, 8 nt upstream of m2_7 seed complementarity, the offset would be +2 nt (8 minus the distance between nucleotides 13 and 7 of the miRNA; 8 – (13–7) = +2). In our naming convention, this site would be designated as m13_20|+2|m2_7, to indicate the 3′ complementarity, offset, and seed complementarity, respectively.

We found that downstream seed complementarity generally resulted in sites that bound better than the analogous 3′-only site acting autonomously, even when that seed complementarity was quite short. To illustrate this finding, representative compound sites are included in Figure 1B. For miR-155, these compound sites had 3’ complementarity from positions 13–22, offsets of +2 nt, and seed complementarity beginning at position 2 and ranging from positions 3 to 8, whereas for miR-124, they had seed complementarity over the same range, but 3’ complementarity from 9–19 and offsets of +3. (Fig 1B). The relative affinities of these compound sites revealed that as few as three nucleotides of contiguous seed complementarity on top of m13_22 3′ complementarity for miR-155 or m9_19 3′ complementarity for miR-124 was sufficient to detectably improve binding (for miR-155, compare m13_22 with m13_22|+2|m2_4; for miR-124, compare m9_19 with m9_19|+3|m2_4). Extending these analyses to additional sites with extended 3’ complementarity and a broad range of offset and seed pairing possibilities confirmed that 3-nt stretches of additional seed complementarity often enhanced affinity, particularly when this involved positions 2 through 4 (Fig S2 and S3). Seed complementarity of more than three nucleotides substantially improved affinity. For example, the affinity of a miR-155 compound site with 10 nucleotides of 3’ complementarity and four nucleotides of seed complementarity (m13_22|+2|m2_5) matched that of a canonical 7mer-A1 site (Fig 1B), and the affinity of the analogous miR-124 site (m9_19|+3|m2_5) exceeded that of a canonical 7mer-A1 site (Fig 1B). For both miRNAs, the degree to which seed complementarity boosted affinity was influenced by offset, with slightly positive offsets resulting in a greater boost to affinity than negative offsets for all possible seed pairing architectures (Fig S2 and S3).

In pursuit of a global analysis that would highlight the respective contributions of 3′ complementarity, seed complementarity, and offset to the relative affinities of compound sites, we fit log-transformed relative *K*_d_ values to a multiplicative model in which a compound site negative log_10_(*K*_d_) was equal to the product of coefficients denoting the contributions of 3′ complementarity, seed complementarity, and offset (A_3’_, B_5′_, C_offset_, respectively). Coefficients were globally fit for the *K*_d_ values of over 5,000 compound sites, such that each coefficient value represented the typical contribution of a particular pattern of 3′ complementarity, pattern of seed complementarity, or offset to overall site *K*_d_ across the many sites in which it appeared. Values of coefficients A, B, and C were each scaled to range from 0 to 1. The model fit our data well (Fig 1D). Although the absolute magnitude of the fitted coefficients had no biochemical significance, comparing relative magnitudes of coefficients predicted the difference in relative affinity. For compound sites of both miRNAs, longer and/or more distal 3′ complementarity typically contributed greater affinity, matching the trends observed for 3′-only sites (Fig 1E, left). In this analysis, 2–3 nucleotides of contiguous seed complementarity provided little-to-no boost to overall site affinity, especially when that seed complementarity occurred outside of the subseed region (nucleotides 2–5 of the miRNA), which is poised to nucleate target pairing (11–17, 22, 33–35). However, 4–5 nucleotides of contiguous seed complementarity typically did provide a detectable boost to compound-site affinity and was perhaps more impactful when it included the subseed region. Sites with at least six nucleotides of contiguous seed complementarity were considered 3′-supplementary sites because the seed complementarity matched that of other canonical sites. For these sites, the relative contribution of different patterns of seed complementarity to overall site affinity matched the relative affinities of the equivalent seed-only sites acting autonomously (Fig 1E, middle). Positive offsets were favored for both miRNAs; offsets of 0–2 were optimal for miR-155, and offsets of 1–4 were optimal for miR-124 (Fig 1E, right). These offset preferences matched those reported for miR-155 and miR-124 in the context of 3′-compensatory sites (27).

### AGO–miR-155 and AGO–miR-124 bind 3′-only sites with high affinity

To measure AGO–miR binding to 3′-only sites in a single target context and determine absolute *K*_d_ values for the binding interactions, we conducted equilibrium binding assays between purified AGO–miR and trace (1–5 pM) radiolabeled target RNA that featured a single target site flanked by poly(U) sequence. Binding reactions were incubated for 4 h at 37°C before filtering through stacked nitrocellulose and nylon membranes that captured bound and free target RNAs, respectively. Fraction bound as a function of AGO–miR concentration was fit to the fractional saturation equation to determine an absolute *K*_d_ value. We also determined affinities for mutant miR-155 and miR-124 sequences in which any A residues in the seed region were mutated to avoid incidental mono- or di-nucleotide pairing between the miRNA seed and poly(U) flanking sequence.

For all four guide sequences (miR-155 (wt), miR-155 (mut), miR-124 (wt), and miR-124 (mut)) *K*_d_ values were determined for multiple 3′-only targets, 7mer-m8 canonical targets, and no-site negative-control targets. The 8-nt 3′-only sites (m13_20 for miR-155, m11_18 for miR-124) did not bind detectably above no-site negative controls (Fig 2A), whereas the 10–11-nt 3′-only sites (m13_22 for miR-155, m9_19 for miR-124) bound with intermediate affinities, with *K*_d_ values ranging between 0.2 and 1 nM (Fig 2A), and the 13–14-nt 3′-only sites (m11_23 for miR-155, m9_22 for miR-124) bound with high affinity, with *K*_d_ values ranging from 20–100 pM— similar to those of the 7mer-m8 sites (Fig 2A). These results concurred with relative affinities measured by AGO-RBNS.

**Figure 2.**
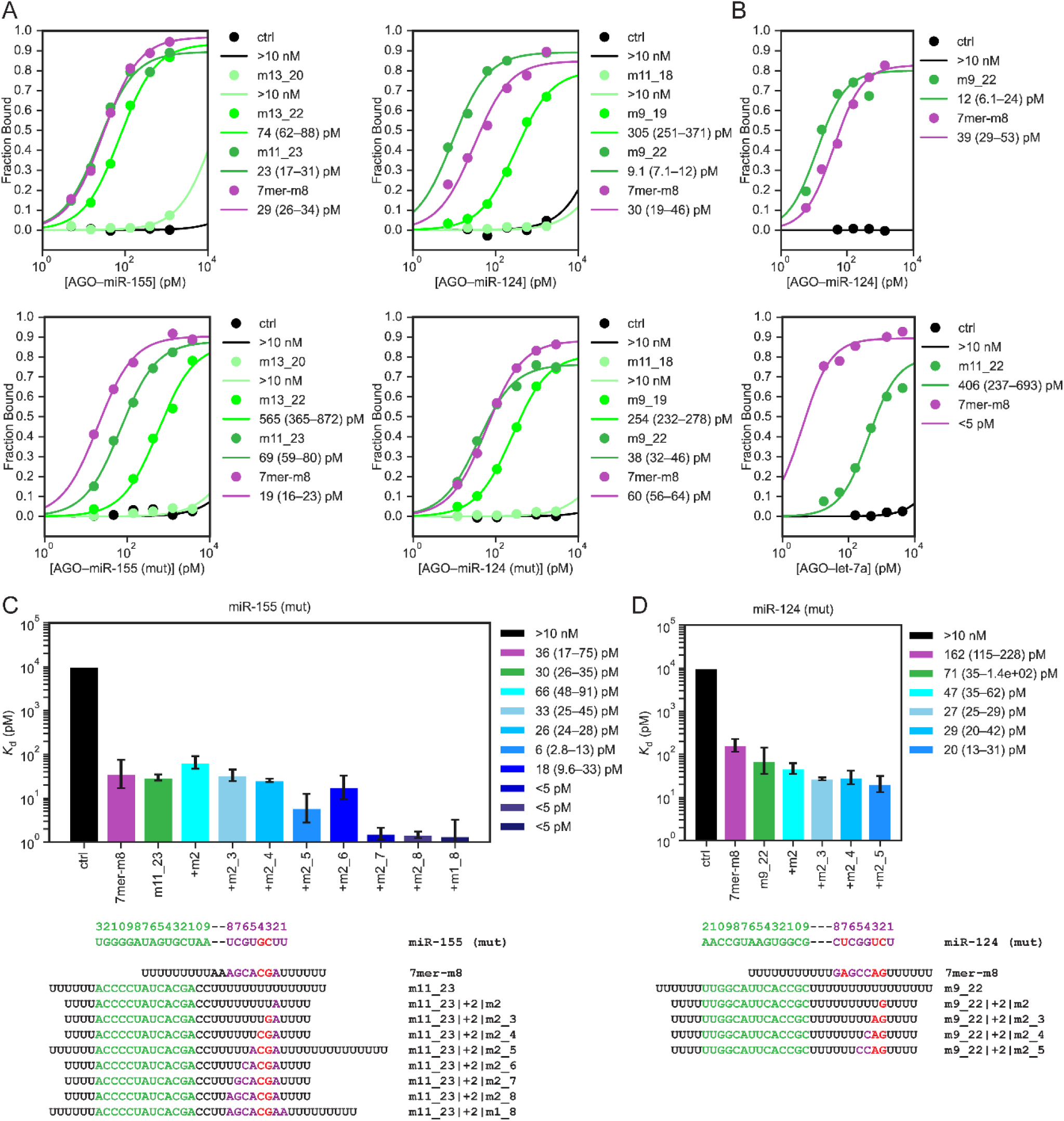
Binding assays confirm high-affinity binding of AGO–miR-155 and AGO–miR-124 to 3′-only sites. (A) Binding curves and absolute *K*_d_ values for AGO–miR-155 and AGO– miR-124 binding to 3′-only sites. Shown are binding curves from single-target equilibrium-binding assays with wild-type guide sequence (top) or mutant guide sequence (bottom) AGO– miR-155 (left) or AGO–miR-124 (right). Targets each contained the indicated single site flanked by poly(U) sequence. Curves were fit to the fractional saturation equation to determine absolute *K*_d_ values reported in the key next to each plot, with 95% confidence interval of each fit. For targets with negligible binding at the highest AGO–miR concentrations, a lower bound of *K*_d_ of 10 nM is reported. For targets that saturated binding at the lowest AGO–miR concentrations, an upper bound of *K*_d_ of 5 pM is reported. (B) Binding curves and absolute *K*_d_ values for AGO– miR-124 (top) and AGO–let-7a (bottom) binding to 3′-only sites. Otherwise, this panel is as in (A). (C) Absolute *K*_d_ values for AGO–miR-155 (mut) binding to compound sites. *K*_d_ values and 95% confidence intervals were determined as in (A) for seed-only (7mer-m8, purple), 3′-only (m11_23, green) and various compound (blue) sites binding to AGO-miR-155 (mut). Sequences of guide and target RNAs are provided below the plot. For targets with negligible binding at the highest AGO–miR concentrations, a lower bound of *K*_d_ of 10 nM is reported and plotted. (D) Absolute *K*_d_ values for AGO–miR-124 (mut) binding to compound sites. Otherwise, this panel is as in (C).

We also measured absolute *K*_d_ values for target RNAs with different combinations of 3′ complementarity and seed complementarity at offsets predicted to be favorable (+2 and +3 nt, for miR-155 and miR-124, respectively). For miR-155 (mut), we designed targets with m11_23 3′ complementarity augmented by 1–8 nucleotides of seed complementarity (m2, m2_3, m2_4, m2_5, m2_6, m2_7, m2_8, m1_8). We found that although 1–3 nucleotides of contiguous seed complementarity did not detectably boost affinity over that of m11_23 3′-only sites (Fig 2C), 4–5 nucleotides of seed complementarity modestly boosted affinity (Fig 2C). Sites with six or more nucleotides of seed complementarity were very strong binders such that fraction bound was already near saturation at the lowest concentrations of AGO–miR (∼5–10 pM), and thus we could report only an upper bound of ∼5 pM for the absolute *K*_d_ of these sites (Fig 2C).

Surprisingly, m2_5 seed complementarity initially appeared to result in a compound site that bound at least as well as one with m2_6 seed complementarity (Fig. 2C), but after controlling more carefully for the length of poly(U) sequence around the site, absolute *K*_d_ values for the two sites were more similar, with m2_6 seed complementarity resulting in slightly stronger binding, as expected (Fig S13).

For miR-124 (mut), we designed an analogous series of targets with compound sites that combined m9_22 3′ complementarity with 1–8 nucleotides of contiguous seed complementarity. Unfortunately, targets with ≥5 nucleotides of seed complementarity were predicted to be highly structured due to unavoidable sequence complementarity between the 3′ region and seed region of miR-124 (mut). Because structure in and around a target site can interfere with AGO– miR binding, particularly if it occludes the portion of the target that pairs to the miRNA seed (23, 28, 31, 36), we limited our investigation to those targets with 1–4 nucleotides of seed complementarity. Again, 1–2 nucleotides of seed complementarity produced no detectable boost to overall site affinity, but 3–4 nucleotides had a modest effect (Fig 2D). As in RBNS experiments, less seed complementarity was necessary to boost compound site affinity for miR-124 than for miR-155 (Fig 2C, 2D), presumably because the seed region (especially the subseed) of miR-124 is more GC-rich.

We also conducted binding assays with AGO–let-7a, which has only marginal enrichment of 3′ *k*-mers in AGO-RBNS analyses (28) and found that let-7a binds its m11_22 3′-only site only weakly (*K*_d_ ∼ 400 pM)—more than 80-fold worse than it binds its 7mer-m8 site (*K*_d_ < 5 pM), consistent with let-7a being substantially less capable of binding 3′-only sites than either miR-155 or miR-124 (Fig 2B).

### Reporter mRNAs containing 3′-only sites are repressed

Having measured strong binding of AGO–miR-155 and AGO–miR-124 to their respective 3′-only sites, we set out to determine whether that strong binding results in effective repression of site-containing reporter mRNAs in mammalian cells. These reporter assays were conducted in a massively parallel format, inserting many different target sites of the two miRNAs into the 3′ UTR of an *EGFP* reporter mRNA in multiple sequence contexts (Fig 3A, S12A). For miR-155, all six canonical sites (6mer-A1, 6mer-m8, 6mer, 7mer-A1, 7mer-m8, 8mer) were represented, each with five different upstream sequences that were designed to avoid 3′ pairing that might supplement those canonical sites (Fig 3A, S12A). In addition to these 30 canonical site architectures, we included the 20 3′-only sites of miR-155 for which we had determined relative affinities by AGO-RBNS. Also included were a selection of 168 compound sites for miR-155, which covered three different patterns of 3′ complementarity (weak (m13_20), intermediate (m13_22), and strong (m11_23)) in combination with 28 different patterns of seed complementarity (all possible patterns of seed complementarity of 2–8 nucleotides) at two different offsets (zero offset and +2 nt offset) (Fig 3A, S12A). Five no-site architectures with no more than three nucleotides of contiguous base pairing to miR-155 were also included as negative controls (Fig 3A, S12A). Each of these 223 miR-155 sites (30 canonical sites, plus 20 3’-only sites, plus 168 compound sites, plus five negative controls) and each of a similarly designed set of 218 miR-124 sites (30 canonical sites, plus 15 3’-only sites, plus 168 compound sites, plus five negative controls), was placed into 20 different sequence contexts, using the same set of 20 50-nt upstream flanking sequences and 20 50-nt downstream flanking sequences for all sites, yielding a total of 8,820 reporter variants in our massively parallel reporter assay (MPRA) (Fig 3A, S12A).

**Figure 3.**
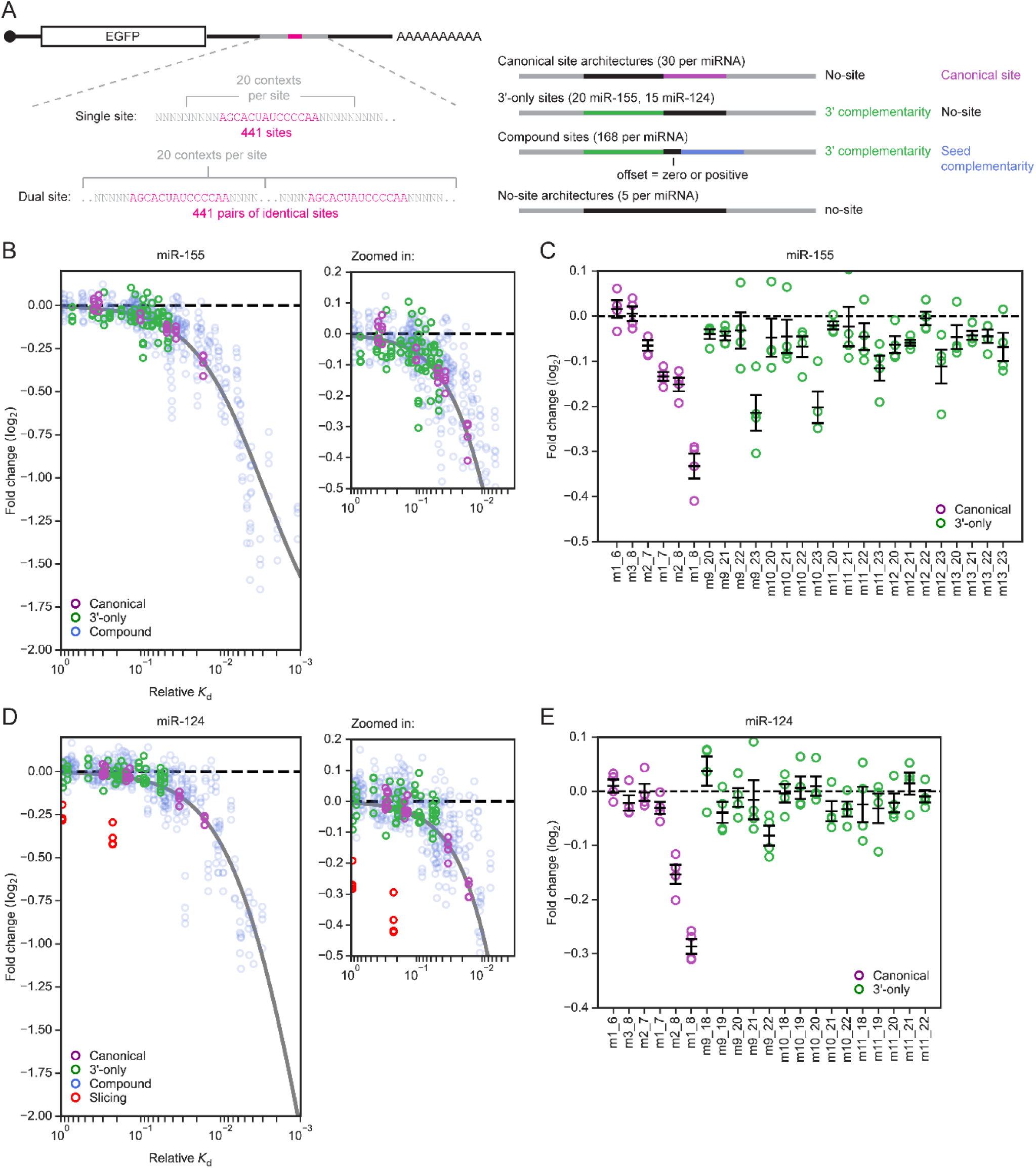
3′-only sites impart repression to reporter mRNAs in F9 cells. (A) Library of variants for massively parallel reporter assay designed to assess the relative repression efficacy of canonical, 3′-only, and compound sites of miR-155 and miR-124, more fully described in Figure S12A. (B) Correspondence between repression of site-containing reporter mRNAs in response to miR-155 transfection and the relative *K*_d_ values of the sites measured by AGO-RBNS (green: 3′-only sites, purple: canonical sites, blue: compound sites). The grey line shows the predicted steady-state repression of sites as a function of relative *K*_d_ from a thermodynamic model of target repression (28). At the right is a zoomed-in view of the top left region of main plot. *N* = 4 replicates per site. (C) Repression efficacy of canonical (purple) and 3′-only (green) sites of miR-155. *N* = 4 replicates per site (error bars, standard error of the mean). (D) Correspondence between repression of site-containing reporter mRNAs in response to miR-124 transfection and the relative *K*_d_ values of sites measured by AGO-RBNS (green: 3′-only sites, purple: canonical sites, blue: compound sites, red: probable slicing sites). Otherwise, this panel is as in (B). Details of how probable slicing sites were identified are provided in Supplementary Information. (E) Repression efficacy of canonical (purple) and 3′-only (green) sites of miR-124. Otherwise, this panel is as in (C).

The reporter library in plasmid form was co-transfected along with guide–passenger duplex for either miR-155, miR-124, or miR-1 into either HeLa or F9 cells. Mock-transfected samples were treated identically except no miRNA duplex was added. After 24 h, cells were harvested, total RNA was extracted, and library mRNAs were reverse transcribed and amplified for high-throughput sequencing. Two replicates were conducted. Reads that exactly matched one of the 8,820 library members were assigned to that member, and for each site, summed counts per million (CPM) values across the 20 contexts in which it appeared were used to calculate fold-change values for each site in the presence of the cognate miRNA compared to mock-transfected samples, normalizing to negative-control values.

We also generated a dual-site library, which was identical to the single-site library except the site region of each mRNA contained two copies of the relevant site, separated by a 20-nt spacer sequence that varied along with the 50-nt flanking sequences that were appended to either side of the site region (Fig S12B). For experiments involving the dual-site libraries, these dual-site libraries were co-transfected together with single-site libraries and miRNA duplexes into F9 cells. Fold-change values for single sites in these experiments correlated extremely well with those of the initial experiments (Fig S7) in which only the single-site library was transfected, and thus these single-site fold-change values were analyzed as replicates, bringing the total number of replicates for the single-site libraries in F9 cells to four for each miRNA. Repression efficacies of sites were generally slightly higher in HeLa cells than in F9 cells, especially for the strongest sites (Fig S5, S6, S7). In both cell lines, repression efficacies of miR-155 sites for miR-155-transfected samples were generally slightly higher than those of miR-124 sites for miR-124-transfected samples, perhaps because either transfection or utilization of miR-155 was more efficient.

For some 3′-only sites, particularly those with stronger relative affinity as measured by AGO-RBNS, repression was detected in cells, thereby demonstrating the intracellular activity of this site type (Fig 3B, 3D). Indeed, for all sites for which we had AGO-RBNS-determined relative affinities, repression efficacy scaled with binding affinity, regardless of whether the site was canonical, 3′-only, or compound (Fig 3B, 3D). Our data fit well to a simple biochemical model that predicts repression as equal to (1+bN) for each site, where N is the fractional AGO–miR occupancy of that site as modelled by the fractional saturation equation, AGO–miR_free_ / (AGO– miR_free_ + relative *K*_d_), b is the unit increase in repression for a unit increase in fractional occupancy, and AGO–miR_free_ is the absolute concentration of AGO–miR-155 or AGO–miR-124 not bound to any RNA (28). Because we could not experimentally determine AGO–miR_free_ or b, these parameters were fit globally (Fig 3B, 3D).

For both miRNAs, many ≥10-nt 3′-only sites repressed at least as well as canonical 6mer sites (Fig 3C, 3E). However, only the few longest and highest-affinity 3′-only sites repressed at least as well as 7-nt canonical sites (7mer-A1 and 7mer-m8) (Fig 3C, 3E). At least 12 nucleotides of contiguous 3′ pairing were required for this strong repression (Fig 3C, 3E). This concurred with our binding affinity data, which showed that 12 nucleotides of contiguous 3′ pairing was required for 3′-only sites to bind as well as top canonical sites.

### Reporter mRNAs containing compound sites are more repressed

Compound sites that had extensive 3′ complementarity and seed complementarity were very efficacious (Fig 4A, 4B). The strongest compound sites of miR-155 had log_2_(fold-change) values less than –2.0, which equated to a >75% reduction in steady-state mRNA abundance, whereas a single 8mer canonical site imparted only a ∼25% reduction (Fig 4A). For sites with the longest 3′ complementarity (m11_23 for miR-155, m9_22 for miR-124) and seed complementarity of at least 5–6 nucleotides, additional seed complementarity did not necessarily increase efficacy, suggesting that these sites were saturated with AGO–miR under our assay conditions (Fig 4A, 4B). For sites with only 8 nucleotides of 3′ complementarity, seed complementarity of 6 nucleotides or more was required to generate detectable repression (Fig 4A, 4B). For sites with intermediate 3′ complementarity of 10–11 nucleotides, seed complementarity of 4–5 nucleotides was sufficient to boost repression beyond that observed for the 3′-only site acting autonomously (Fig 4A, 4B). For sites with the longest 3′ complementarity, additional seed complementarity of as few as 2–3 nucleotides was sufficient to boost repression beyond that observed for the 3′-only site acting autonomously (Fig 4A, 4B). Note that our ability to observe the activity of as few as 2–3 nucleotides of additional seed complementarity in the reporter assay but not in AGO-RBNS was attributed to the inclusion of compound sites with particularly extensive 3’ complementarity in the MPRA, whereas our AGO-RBNS analysis was not powered to derive relative *K*_d_ values for these particularly long compound sites because they were insufficiently abundant in the input library.

**Figure 4.**
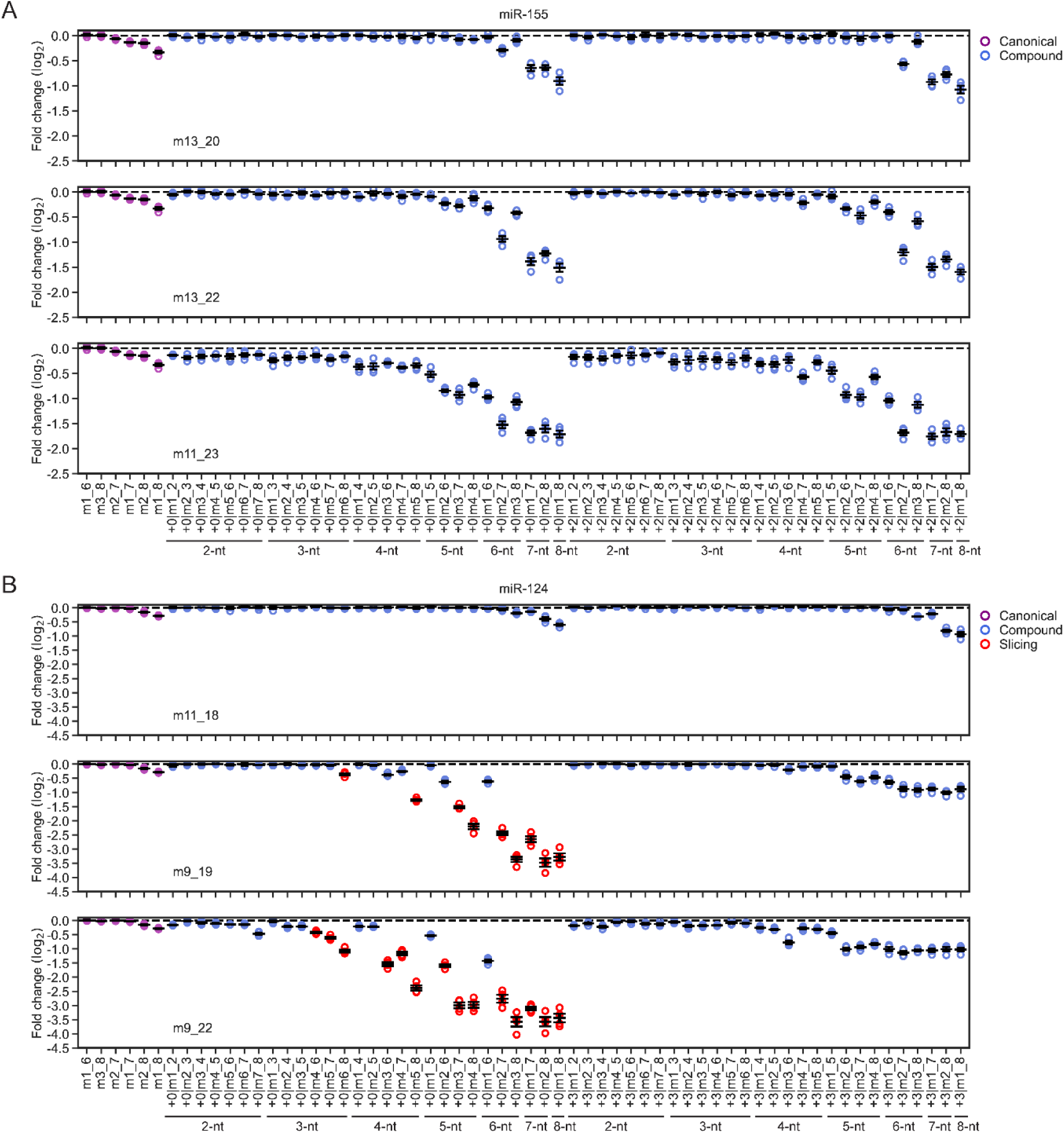
Repression efficacy of compound sites depends on 3′ complementarity, seed complementarity, and offset. (A) Repression efficacy of reporter mRNAs containing miR-155 target sites (purple: canonical sites, blue: compound sites) in response to miR-155 transfection. Top, middle, and bottom plots show compound sites with m13_20, m13_22, and m11_23 3′ complementarity, respectively. As an example of site nomenclature, +0|m1_3 indicates that in addition to the 3′ complementarity, the site featured m1_3 seed complementarity at an offset of +0. *N* = 4 replicates per site (error bars, standard error of the mean). (B) Repression efficacy of reporter RNAs containing miR-124 target sites (purple: canonical sites, blue: compound sites, red: probable slicing sites). Otherwise, this panel is as in (A). Details of how probable slicing sites were identified are provided in Supplementary Information.

For compound sites of miR-155 that did not appear saturated, repression efficacy was better for positive offset (+2) compared to no-offset (+0) versions of equivalent sites (Fig 4A), as observed previously for 3’-supplementary sites (27). Compound sites of miR-124 with intermediate (m9_19) or strong (m9_22) 3′ complementarity and additional seed complementarity at zero offset were expected to be suitable substrates for slicing by AGO2– miR-124, especially when seed complementarity extended to position 8 and thereby created perfect complementarity through the seed and central regions (Fig 4B). Therefore, we could not compare positive-offset and zero-offset versions of these compound sites. Nevertheless, for sites with the weak (m13_20) pattern of 3′ complementary, the expected preference for positive offset was observed.

Slicing of the miR-124 reporter was inferred by increased repression observed with an offset of +0 nt compared to that observed for the analogous site with an offset of +3 nt, because the positive offset would correspond to a 3-nt bulge between positions 8 and 9 which is poorly tolerated for slicing (23). Detection of reporter sites that were sliced provided an opportunity to compare slicing results that we observed in cells for miR-124 to those previously observed in vitro for other miRNAs (23). In the context of intermediate-strength (m9_19) 3′ complementarity, m6_8 or m5_8 seed complementarity (creating contiguous complementarity from positions 6–19 and 5–19 respectively) was sufficient to create a weak slicing site (Fig 4B). If complementarity through nucleotides 3–7 of the seed was intact, a single mismatch at position g8 was well-tolerated (Fig 4B). In the context of strong (m9_22) 3′ complementarity, a double mismatch at positions 7–8 was insufficient to completely abrogate slicing, and as little as m7_8 seed complementarity (creating contiguous complementarity from positions 7–22) was sufficient to generate at least some slicing, demonstrating that miR-124 had relaxed seed complementarity requirements for slicing in the context of more extensive 3′ complementarity (Fig 4B). These results were consistent with those observed in vitro for miR-21 and let-7a, for which partial seed mismatches do not completely abrogate slicing as long as perfect central and 3′ complementarity are preserved (23).

Overall, long stretches of contiguous 3′-complementarity imparted repression to reporter mRNAs according to their measured binding affinity, whether or not they acted autonomously as 3′-only sites, or were supplemented by additional base pairs to the seed region. Eight nucleotides of 3′ complementarity was insufficient for detectable repression, unless supplemented by canonical 6–8-nt seed complementarity. 10–11 nucleotides of contiguous 3′ complementarity imparted detectable but modest repression similar to that imparted by a 6-nt canonical site, but 4–5 nucleotides of additional seed complementarity could boost that repression and create highly efficacious sites. 12 or more nucleotides of contiguous 3′ complementarity imparted effective repression similar to that imparted by a 7-nt canonical site, and as few as 2–3 nucleotides of additional seed complementarity further boosted that repression, creating sites that repressed better than 8mer canonical sites.

### 3′-only sites have slower association rates than canonical sites

The pairing of miRNAs to extended target sites is thought to be an ordered process. Pairing nucleates at the subseed and then propagates to the remainder of seed region (22, 42–44, 46). Pairing to the full seed is also proposed to cause conformational changes in AGO that open the 3′-supplementary chamber to make pairing to the 3’ region more accessible (46). Pre-organization of the subseed into a helical conformation results in rapid association rates for nucleation—often approaching diffusion-limited bimolecular association (21–23, 34, 42–44), raising the question of how rapidly AGO–miRs might associate with 3′-only sites, which lack seed pairing and consequently, the benefits of both the pre-organized guide and the widened supplementary chamber proposed to make 3′ pairing more accessible (46). Previous analyses using AGO–let-7 found that this miRNA had an elementary rate constant for target association (*k*_on_) of 3.6 ± 0.2 x10^7^ M^−1^s^−1^ for the 3′-only site m11_21, compared to 2.4 ± 0.1 x10^8^ M^−1^s^−1^ for a canonical 8mer site (22). However, let-7 is not prone to undergo stable 3’-only pairing (Fig 2B), and thus we set out to measure the *k*_on_ for a 3’-only site of miR-155, which underwent stable 3′-only pairing in AGO-RBNS and single-target equilibrium-binding assays (Figs 1B,C, and 2A).

To measure association rates of AGO–miRs with different target sites, we conducted binding reactions with single-site target RNAs wherein the site was flanked by poly(U) sequence (Fig 5A). To increase the number of sites tested at once, we used targets with different lengths of flanking poly(U) sequence, such that when using a single binding reaction containing multiple targets, the amount of each target that was bound at each timepoint could be resolved on a denaturing gel (Fig 5A). For each target, the bound fraction was fit to the integrated rate equation for pseudo first-order bimolecular association, and a *k*_on_ value was determined (Fig 5A). Experiments were repeated with two other concentrations of AGO–miR, and *k*_on_ values from these replicates were generally very similar (Fig 5B). Experiments were also performed for two different guide sequences: miR-155 (wt) and miR-155 (mut) (Fig 5B).

**Figure 5.**
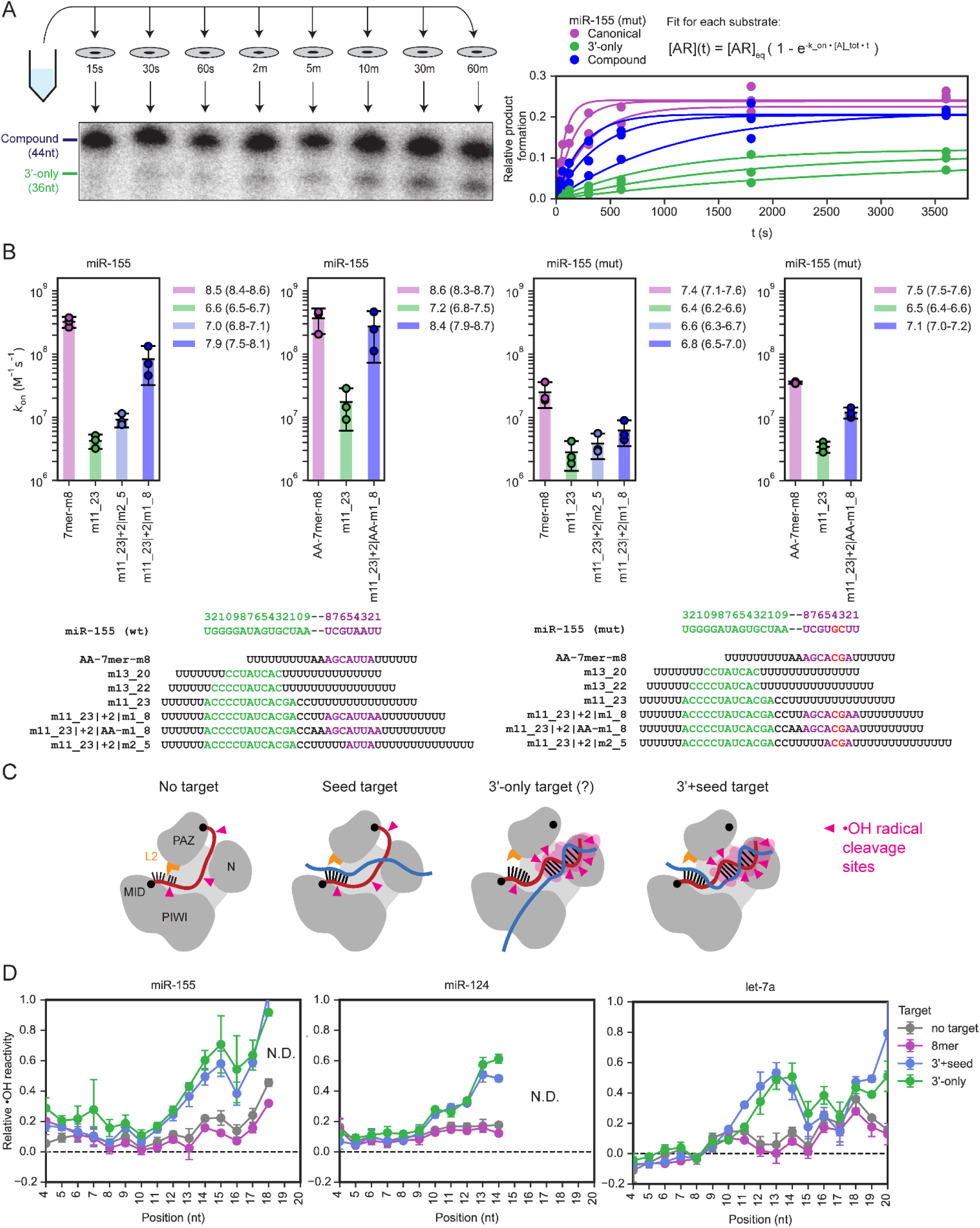
AGO–miR-155 and AGO–miR-124 engage 3′-only sites by an alternative seedless mechanism. (A) Schematic and examples of assays measuring pseudo-first-order association rates. (B) Association rates for wild-type and mutant miR-155 binding to canonical, compound, and 3′-only sites. Plotted are *k*_on_ values for wild-type AGO–miR-155 (panels 1–2) and AGO–miR-155 (mut) (panels 3–4) binding to targets with either the canonical 7mer-m8 site (purple), a 3′-only site (green), or one or two compound sites [light blue: compound site with partial (m2_5) seed complementarity, dark blue: compound site with full (m1_8) seed complementarity]. Panels 1,3 feature a first set of targets; panels 2,4 feature a second set of targets with the dinucleotide upstream of seed pairing altered to reduce structuredness. *N* = 3 replicates (error bars, 95% confidence interval). Sequences of guide and target RNAs are depicted below plot. (C) Schematic of hydroxyl radical probing assays. (D) Backbone accessibility patterns observed for miRNAs of AGO–miRs bound to either no target or to targets with different types of sites. Plotted are relative backbone accessibilities at each informative position along the miRNA guide for AGO–miRs pre-saturated with either buffer-only negative control (grey), a target with an 8mer canonical site (purple), a target with a 3′-only site (green), or a target with extensive 3′ complementarity and full m1_8 seed complementarity (3′+seed, blue). Results are shown for miR-155 (left), miR-124 (middle), and let-7a (right). *N* = 2 replicates for miR-155 and let-7a (error bars, range). *N* = 3 replicates for miR-124 (error bars, 95% confidence interval).

Association rates of 7mer-m8 canonical sites were fast, with *k*_on_ values of 10^8.5–8.6^ M^−1^s^−1^ and 10^7.5–7.6^ M^−1^s^−1^ for miR-155 (wt) and miR-155 (mut), respectively (Fig 5B). These values were within range of those reported previously for seed-based target engagement of other guides (21–23, 34, 42–44). For m11_23 3′-only sites, *k*_on_ values were 10^6.5–7.2^ M^−1^s^−1^ and 10^6.4–6.6^M^−1^s^−1^ for miR-155 (wt) and miR-155 (mut), respectively (Fig 5B), similar to the previously reported *k*_on_ value of the let-7 m11_21 site (22). These association rates for 3′-only sites were 10–100-fold slower than those of 7mer-m8 sites, consistent with the hypothesis that seedless engagement with 3′-only sites would be slower than seed-based engagement with canonical sites. These slower association kinetics of 3’-only sites implied correspondingly slower dissociation of 3′-only sites, compared to seed-matched sites of the same affinity, which explained why 3’-only site binding appeared slightly stronger in our single-target equilibrium binding assays than in our AGO-RBNS experiments. The shorter incubation time for our AGO-RBNS experiments (1–2 h compared to 4 h for our single-target assays) was likely insufficient for these sites with slower dissociation rates to approach binding equilibrium, and thus our AGO-RBNS experiments slightly under-estimated the relative affinity of the strongest 3’-only sites.

Compound sites with both 3′ complementarity and seed complementarity were expected to have fast association rates, resembling those of 7–8-nt canonical sites, because their seed pairing should allow them to engage AGO–miRs through the fast, seed-based mechanism. However, for compound sites with m11_23 3′ complementarity and full m1_8 seed complementarity (m11_23|+2|m1_8 sites, also classified as 3’-supplementary sites), we initially observed association rates falling between those of 3′-only sites and 7mer-m8 sites (Fig 5B, panels 1,3). These *k*_on_ values were 10^7.9^ M^−1^s^−1^ and 10^6.8^ M^−1^s^−1^ for miR-155 (wt) and miR-155 (mut), respectively. We investigated the possibility that RNA structure might obstruct accessibility of these compound sites, as structure in targets, particularly structure involving the portion of a target that pairs with the miRNA seed, can substantially slow AGO–miR association (23, 34, 42, 45). Accordingly, we predicted the secondary structures of these target RNAs, recording for each base along the target the predicted probability of being involved in a base pair, as an indicator of structural occlusion. For both guides, the targets with 3′-supplementary sites (m11_23|+2|m1_8) appeared to be more structurally occluded than those with 7mer-m8 sites, especially in the seed region (Fig S8). Some of this occlusive pairing depended on the dinucleotide immediately upstream of the m1_8 seed pairing; changing this UU dinucleotide to AA in targets with the compound sites resulted in targets predicted to be less structured, particularly for the one with the compound site for miR-155 (wt) (Fig S8). Indeed, this modified 3′-supplementary site for miR-155 (wt) (designated m11_23|+2|AA-m1_8) had a *k*_on_ value of 10^8.4^ M^−1^s^−1^)—nearly identical to that of the 7mer-m8 site (Fig 5B, panel 2 of 4). The target with the modified 3′-supplementary site for miR-155 (mut), which was still predicted to be slightly more structured than that with the 7mer-m8 site, had a *k*_on_ value of 10^7.1^ M^−1^s^−1^, which was also faster than the original target (10^6.8^ M^−1^s^−1^) but not quite as fast as that with the 7mer-m8 canonical target (10^7.4-7.5^ M^−1^s^−1^) (Fig 5B, panel 4 of 4). Thus, on the whole, after accounting for target structure, compound sites with m1_8 seed pairing had association rates indistinguishable from those of 7–8-nt canonical sites, consistent with previous reports for other guides (23).

More intriguing was the case of compound sites with 3′ complementarity and only partial seed complementarity (also classified as 3’-compensatory sites). We focused on m11_23|+2|m2_5 sites to miR-155 (wt) and miR-155 (mut), which featured complementarity to seed nucleotides 2–5 in addition to 3′ complementarity to nucleotides 11–23. This pattern of seed complementarity was chosen because the subseed (guide nucleotides 2–5) is the region of the miRNA thought to first contact incoming targets and is thus primarily responsible for the fast association rates of AGO–miRs with seed-matched sites (22, 23, 43, 44). We found that these m11_23|+2|m2_5 compound sites had *k*_on_ values of 10^6.6^ M^−1^s^−1^ and 10^7.0^ M^−1^s^−1^ for miR-155 (wt) and miR-155 (mut), respectively, which were 10–40-fold slower than those of the 7mer-m8 sites. Indeed, these values for m11_23|+2|m2_5 compound sites were only 2–3-fold faster than the those of m11_23 3’-only sites (Fig 5B, panels 1,3). Unlike compound targets with m1_8 seed complementarity, these compound targets with m2_5 seed complementarity were not predicted to be significantly more structured than 7mer-m8 targets (Fig S8), arguing against structural occlusion as an explanation for their surprisingly slow association rates. Others have reported that mismatches at positions 6–8 in the context of otherwise perfectly complementary targets minimally affect association rate for miR-21 but substantially decreases association rate for let-7a (23). Thus, our results show that miR-155 (wt) and miR-155 (mut) behave more like let-7a, in that pairing to the distal part of the seed region appears to make a substantial contribution to the fast association of sites that contain seed pairing.

### Association with 3′-only sites involves release of the miRNA 3′ end from the PAZ domain of AGO

To characterize the conformation of AGO–miR bound to 3′-only sites, we conducted chemical probing experiments to measure changes in the conformation of the miRNA guide backbone upon binding to 3′-only sites, as well as canonical sites and 3′-supplementary sites. Purified AGO–miR, with radiolabeled guide, was incubated with a saturating target RNA, and then Fe(II)–EDTA was added to generate hydroxyl radicals, which cleave RNA at solvent-accessible backbone positions, with the degree of cleavage corresponding to the relative solvent accessibility of the backbone ribose (Fig 5C). After quenching, the ensemble of cleavage products was resolved on denaturing gels, and the relative abundance of each radiolabeled cleavage product, which indicated the solvent accessibility of the respective guide RNA backbone position, was quantified.

Although relative •OH reactivity could not be determined for every position of all guides, clear patterns were observed (Fig 5D). First, for all three guide sequences (miR-155, miR-124, let-7a), binding to only the seed had minimal impact on the pattern of backbone accessibility, with the results observed for AGO–miR incubated with a target with a canonical 8mer site closely matching those observed for AGO–miR incubated with no target (Fig 5D). These results concurred with probing results for other miRNAs (34) and expectations based on crystal structures, which show minimal changes in guide-RNA solvent accessibility upon pairing to the seed region (46). In contrast, for targets containing a compound site with extensive 3′ complementarity, increased backbone accessibility of the guide 3′ region was observed (Fig 5D), as expected, because pairing beyond nucleotides 17–18 of the guide disrupts backbone contacts to PAZ and other regions of the protein (47). Moreover, because the guide 3’ region and target wrap around each other to form the 3′ helix, some periodicity in guide backbone accessibility was also expected when pairing to these compound sites, due to decreased accessibility of the backbone positions facing inwards towards the wall of the 3’-supplementary chamber (34, 47). Indeed, for both miRNAs for which sufficient probing data could be acquired, a dip in accessibility centering on position 16 was observed (Fig 5D). Very similar patterns of backbone accessibility were observed when pairing to 3′-only sites, which featured 3′ complementarity identical to that of the compound sites but no seed complementarity (Fig 5D). These results indicate that when bound to 3′-only sites, the 3′ end of the guide is released from PAZ and forms a 3′ helix indistinguishable from that of a 3’-supplementary site.

### 3′-only sites and canonical sites bind AGO–miR with divergent energetics

AGO re-shapes the energetics of target binding for the miRNA guide (22, 23, 27, 28). For instance, for seed-matched sites, the pre-organization of the miRNA seed pre-pays the entropic cost of duplex formation, and thus experimental ΔG° values are more negative than predicted for the equivalent helix forming between two RNA molecules free in solution (22, 23, 27, 28). On the other hand, segments of 3’ pairing within extensively complementary sites are typically less stable than predicted for equivalent RNA molecules free in solution (22, 27). To extend this type of analysis to the 3’-only sites of miR-155 and miR-124, we transformed relative *K*_d_ values from AGO-RBNS to experimental ΔG° values, using the relationship of ΔG° = RTln(*K*_d_) for 6–7-nt canonical sites (not including 6mer-A1, 7mer-A1 or 8mer sites, which have a non-pairing interaction at position 1), 3′-only sites, and compound sites, and compared those to ΔG° values predicted for the equivalent pairing calculated using nearest-neighbor rules for RNA molecules free in solution (48). As expected, 6–7-nt canonical sites had more negative ΔG° values in the context of AGO than predicted for equivalent pairing in solution (Fig 6). In contrast, 3′-only sites and compound sites with 3′ complementarity and m2_4 or m2_7 seed complementarity all had less negative ΔG° than predicted for equivalent pairing in solution (Fig 6). These results supported the idea that AGO imparts a substantial penalty for pairing to the 3’ region, which largely offsets the benefit of 3’ pairing (27, 49). The sources of this inferred penalty are the multiple favorable contacts between AGO and its guide, including binding of the guide 3′ terminus by the PAZ domain of AGO, which must all be disrupted for 3′ pairing to propagate to the 3′ end of the guide (47, 49). The slopes of the lines of best fit for 3′-only and compound sites were more gradual than that of seed sites, indicating that individual 3′ region base pairs contribute less to overall site affinity than individual seed region base pairs (Fig 6). Compound sites with m2_7 seed pairing had the most gradual slope, although we suspect this result is attributable to the possibility that binding to compound sites with extensive 3′ complementarity and m2_7 seed complementarity had not reached equilibrium, due to their very slow dissociation rates, and thus AGO-RBNS relative *K*_d_ values underestimated their affinity.

**Figure 6.**
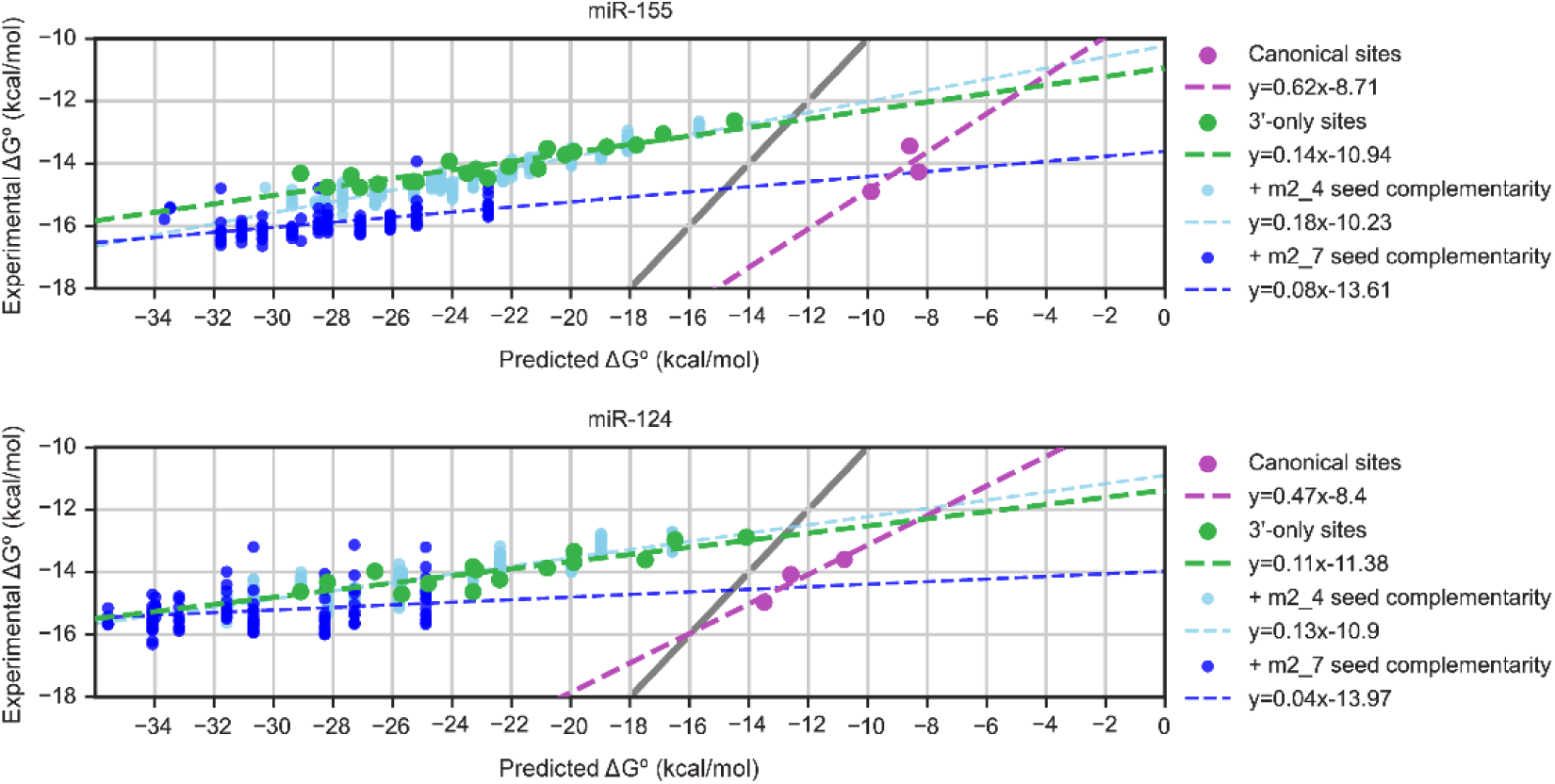
3′-only sites and canonical sites bind with divergent energetics. ΔG° values calculated from experimentally determined relative *K*_d_ values are plotted as a function of the ΔG° values predicted for the analogous pairing in solution for miR-155 (top) and miR-124 (bottom). Results are shown for canonical sites (purple), 3′-only sites (green), compound sites with m2_4 seed complementarity (light blue), and compound sites with m2_7 seed complementarity (dark blue). Grey line shows *y* = *x*.

### 3′-only sites are a minor but potentially important part of the targetome of most conserved miRNAs

Because 3′-only sites involve more contiguous complementarity to the miRNA than canonical sites, they are expected to be rarer in endogenous 3′ UTRs. For example, a 10-nt 3′-only site, expected to occur every 4^10^ nucleotides by chance, is expected to be about 250-fold less common than a 6-nt canonical site, expected to occur every 4^6^ nucleotides by chance. To further examine the relative abundance of 3’-only sites, we searched endogenous human 3′ UTR sequences for predicted target sites of the 183 human miRNAs from the 108 seed families conserved to zebrafish (Fig S9). We first searched for 6mer, 7mer-A1, 7mer-m8 and 8mer canonical sites (with or without 3′-supplementary pairing). We excluded offset 6mer sites (6mer-A1, 6mer-m8) from this analysis because although abundant, they are only marginally, if at all, effective in cells. We also searched for sites that featured at least ten nucleotides of contiguous 3′ complementarity. If seed complementarity of 3–5 nucleotides was present at an offset of between –4 and +6, the site was counted as a “3′+microseed” compound site. If no seed complementarity of three or more nucleotides was present at an offset of between –4 and +6, the site was counted as a 3′-only site.

We focused on 3′-only and 3′+microseed sites with ≥10 nucleotides of contiguous 3′-complementarity because 10-nt 3′-only sites of miR-155 and miR-124 generally bound and repressed about as well as 6mer canonical sites. We found substantial inter-miRNA differences in the relative abundance of canonical versus 3′-only sites in 3′ UTRs (Fig S9A, S9B). Whereas for some miRNAs, almost 100% of assignable sites were canonical sites (e.g., miR-375, 99.96%), for others, less than 90% were canonical (e.g., miR-191, 85.78%). For some miRNAs, 3′-only sites (e.g., miR-126, 9.26%) and 3′+microseed sites (e.g., miR-126, 2.31%) together constituted an appreciable portion of the predicted targetome, whereas for other miRNAs, 3′-only sites were very lowly abundant (e.g., miR-137, 0.05% 3′-only, 0.03% 3′+microseed).The median values for broadly conserved miRNAs were 60 3′-only sites (range: 0–277) (Fig S9B) and 28 3′+microseed sites (range: 0-167) (Fig S9C), compared to 9,573 canonical sites (range: 726-20,873) (Fig S9A). In sum, the median proportion of the predicted targetome contributed by 3′-only sites was about 0.6%, with another ∼0.3% contributed by sites with equivalent 3′ complementarity supplemented by 3–5-nt seed complementarity. Thus, these sites appear to constitute a proportion of the endogenous miRNA targetome resembling that estimated for 3′-compensatory sites (26). Individual 3′-compensatory sites can mediate important miRNA–target interactions (25, 50), underlining the fact that comparative rarity of a particular site type does not preclude its functional importance. The fact that strong binding to 3′-only sites occurs for some miRNAs but not others casts additional uncertainty as to which endogenous 3′-only sites might be functional or consequential. Thus, our estimate of the proportion of the targetome contributed by endogenous 3′-only sites might best be considered as an upper bound.

## DISCUSSION

### Seedless engagement with 3′-only sites

The affinity and efficacy of 3’-only sites demonstrates that 3′ pairing can initiate between a guide RNA and a target site for which there is no seed pairing. This result contrasts with the proposal that the guide RNA 3’ region is inaccessible and improperly oriented for base pairing prior to conformational changes that are brought about by seed pairing (46). In this previous proposal, which is based on crystal structures of AGO2–miRs with-and-without seed-paired target RNA, seed pairing promotes conformational shifts both in the protein and the guide RNA, which widen the channel between the PAZ and the N domains to allow access to the miRNA 3’ region and orient the bases for more facile pairing. Based on our finding that 3’ pairing can initiate at sites without seed pairing and form a 3’ helix indistinguishable from that of a 3’-supplementary site (Fig 5D), we suggest that in the absence of target, AGO–miR populates multiple conformational states, including the state that is more open and optimally oriented for 3′ pairing, and that it dynamically interconverts between these states. Then, upon seed pairing, it more frequently populates that more open, more optimally oriented state. In support of this structurally dynamic model, the 3’ region of the guide is disordered and thus poorly defined in the crystal structure of AGO2 associated with a single guide RNA but no target—implying multiple conformations of the 3’ region (15). Moreover, studies using single-molecule Förster resonance energy transfer and cryogenic electron microscopy show that AGO2 is conformationally flexible (32, 51), consistent with the idea that the crystal structures that suggest inaccessibility of the 3’ region prior to seed pairing might not capture the more open conformations that enable 3’ pairing.

In this dynamic model, one explanation for the different binding affinities of 3′-only sites for different miRNAs is that some guides might have 3′ regions that are more frequently accessible or suitably organized for 3′ pairing. Indeed, the 3′ region of the guide is poorly defined in the crystal structure of AGO2 associated with multiple guide RNAs but no target, implying that different guide RNAs have different conformations of their 3’ regions (17), some of which may be more frequently accessible and better oriented for target pairing, ultimately resulting in different association rates for binding to 3′-only sites for different guides.

One conformational change that might be particularly consequential for nucleation or propagation of 3’ pairing is the release of the guide-RNA 3’ end from the PAZ domain. Dissociation of the 3′-end–PAZ interaction, even in the absence of bound target, might make the 3’ region transiently accessible, with the fraction of molecules populating this dissociated state affecting the association rate. We used periodate chemistry to probe the degree of 3′-end–PAZ dissociation for purified AGO–let-7a, –miR-196a, –miR-7, and –dre-miR-430a and found evidence for appreciable dissociation in the absence of bound target for all 4 guides (Fig S10). Although we did not observe significant inter-guide differences in PAZ dissociation, others have measured binding between purified AGO2 PAZ domain and mono- and poly-nucleotides and observed nucleotide identity preferences (52, 53). If the 3′ ends of certain guides are less strongly bound by PAZ, then those guides would be expected to more frequently populate the dissociated state, perhaps resulting in faster association with 3′-only targets, akin to (and perhaps in combination with) inter-guide differences in 3′ region accessibility.

Taken together, conformational states that impart accessibility or favorable nucleotide orientation within the 3′ region, or dissociation of the 3′-end–PAZ interaction, are all expected to enhance the association rate of 3’-only sites, with the partial occupancy of these states explaining the slower association rates of 3′-only compared to seed-matched sites, and differential population of these states for different 3’ sequences explaining inter-guide differences in 3′-only site binding affinity. In addition to inter-guide differences in 3′-only site association rate, we expect inter-guide differences in 3′-only site dissociation rate, with more stably paired 3′ region sequences resulting in slower dissociation rates and concomitantly higher affinity for 3′-only sites for that guide.

### The contribution of 3′-only sites to endogenous miRNA-mediated regulation

We estimate that 0.5–1% of the endogenous targetome of most broadly conserved miRNAs consists of 3′-only sites with at least 10 nucleotides of contiguous 3′ pairing. However, we do not yet know which of these predicted miRNA–target interactions are consequential. In addition, the comparative rarity of 3′-only sites makes it difficult to quantify the degree to which they impart repression to endogenous site-containing mRNAs. For example, analysis of mRNA-seq data from HeLa cells that had been transfected with miR-155 or miR-124 (28) identified only one and two transcripts that contained a ≥10-nt 3′-only site to these respective miRNAs, after filtering for transcripts with suitable read coverage and a single unambiguously assignable dominant 3’ UTR isoform containing a single site to the relevant miRNA. This was insufficient to compare and statistically test the changes observed for site-containing transcripts compared to those of length-matched no-site control transcripts. Using a more permissive length cutoff of ≥8 nucleotides resulted in numbers sufficient to conduct a statistical analysis for each of the 16 miRNAs that had been transfected (28), but no statistically significant repression was observed, despite a clear signal for canonical sites (Fig S11A). This result was expected when considering that 8 nucleotides of contiguous 3’ complementarity was insufficient for binding and repression by either miR-155 or miR-124. By aggregating 3’-only sites of all 16 miRNAs, we were able to gain enough examples of ≥10-nt 3’-only sites (77) to perform the analysis but still did not observe a signal for measurable repression, presumably because only some miRNAs engage in 3’-only targeting. We suspect that detecting repression from endogenous 3’-only sites will depend on the ability to determine the cohort of miRNAs that engage in 3’-only targeting.

Analysis of the degree to which endogenous 3′-only sites are evolutionarily conserved is also hampered by their comparative rarity. To rigorously quantify the conservation of a particular site architecture, branch-length scores for site sequences across a multiple sequence alignment are calculated and then compared to scores for a cohort of control sequences that speak to the probability that the site sequence is conserved by chance (26). Once again, the small numbers of 3′-only sites of miR-155 and miR-124 was expected to be insufficient for this purpose, and aggregating sites across miRNAs, as done previously for 3’-supplementary sites (26), was expected to be unhelpful because only a subset of miRNAs have functional 3’-only sites that might be conserved. Thus, as for analyses of repression mediated by endogenous 3’-only sites, more selective aggregation appears necessary to illuminate the degree to which endogenous functional 3’-only sites are evolutionarily conserved.

In addition to their differences in relative target abundance, 3′-only and seed-based targeting also differed with respect to their binding kinetics. 3′-only sites had at least 10-fold slower association rates than seed-matched sites, and thus for seed-matched and 3′-only sites of equivalent affinity, the dissociation rate of the 3′-only site is presumably also at least 10-fold slower. These slower rates will increase the time to reach binding equilibrium for 3′-only sites as a group compared to seed-matched sites, implying a delay to reach steady-state repression after induction of a miRNA.

In sum, 3′-only sites of miR-155 and miR-124 that feature extensive 3′ complementarity (10–12 nucleotides or more) can bind with high affinity and impart effective repression to mRNAs in cells, despite their lack of seed pairing, making 3′-only sites a minor but potentially important part of the targetome of certain miRNAs. The next steps will be to determine for which miRNAs the 3′-only targeting mode is operative and why, and to characterize the contribution of 3′-only targeting to endogenous miRNA-mediated regulation.

## MATERIALS AND METHODS

### Purification of AGO–miRs

Plasmid encoding either 3X-FLAG-tagged human AGO2 (pcDNA3.3-3xFLAG-AGO2, Addgene, 136687) (28), 3X-FLAG-SUMO^Eu1^-tagged human AGO2 (pcDNA3.3-3xFLAG-SUMO^Eu1^-AGO2, Addgene, 231372) (43) or 3X-FLAG-SUMO^Eu1^-tagged human AGO2^D669A^ (pcDNA3.3-3xFLAG-SUMO^Eu1^-AGO2(D669A), Addgene, 231371) (43) were transiently transfected into either adherent HEK293T cells or suspension Expi293F cells (Gibco, A14527), along with pMaxGFP (Lonza, discontinued) to assess transfection efficiency. Transfected cells were lysed and AGO-overexpressing S100 cytoplasmic extracts were prepared. For preparation of AGO–miR-155 and AGO–miR-124 for AGO-RBNS experiments, AGO–miR-124 for hydroxyl radical probing experiments, and AGO–miRs for periodate probing experiments, eight 15-cm plates of adherent cells were transfected, lysed, and used to prepare S100 extract, as described (28). For all other AGO–miR preps, 250 mL of suspension cells were transfected, lysed, and used to prepare S100 extract, as described (43).

Preparation of purified AGO loaded with an individual miRNA of interest from AGO-overexpressing S100 extracts was accomplished by a capture–competitor workflow (49), modified as described (43, 50). In brief, synthetic miRNA guide–passenger duplex was added to an aliquot of S100 extract (amounts of extract and duplex varied based on the amount of purified AGO–miR that was required) and incubated with agitation for 1–2 h before addition to streptavidin beads (Dynabeads MyOne Streptavidin C1, Invitrogen, 65002) that had been pre-coated with a 3′-biotinylated capture 2ʹ-O-methylated RNA oligonucleotide (IDT) that contained an 8mer site to the miRNA being loaded. After capture of AGO loaded with the guide of interest, saturating 3′-biotinylated competitor DNA oligonucleotide (IDT) was added, which was fully complementary to the capture oligonucleotide and competed AGO–miR off of the capture beads. The eluent from this competition step was then incubated with magnetic anti-FLAG beads (Anti-FLAG M2 magnetic beads, Milipore, M8823) to affinity purify tagged AGO–miR. Elution from the anti-FLAG beads was accomplished either by competition with free 3X-FLAG peptide (Sigma-Aldrich, F4799) for preps that began with adherent cells (28), or by proteolytic cleavage using SENP^EuB^ protease (purified as described (51)) for preps that began with suspension cells (43). Final eluent was flash frozen in liquid nitrogen in 10 μL single-use aliquots, which were stored at –80°C. AGO–miR preps were quantified at time-of-use by saturation filter binding to a target RNA containing a canonical 7mer-m8 site, as described in supplementary methods.

For AGO–miR preps intended for hydroxyl radical and periodate probing experiments, this workflow was conducted with guide RNAs that had been radiolabeled (as described in supplementary methods), glycerol was omitted from all buffers throughout the purification process to avoid glycerol in the AGO–miR prep reacting with Fenton’s reagent or periodate during probing (50), and the AGO2^D669A^ active-site mutant was employed in place of wild type AGO2 to prevent slicing of target RNAs that were fully complementary to the loaded guide.

### AGO-RBNS

AGO-RBNS was conducted as described (28), with two modifications. First, instead of using 100 nM of random-sequence molecules, the library contained 50 nM of random-sequence molecules combined with 50 nM of programmed molecules that each had 8 nucleotides of complementarity to the 3′ region of the cognate miRNA. Second, libraries were pre-incubated with a 1.1-fold molar excess of a blocking DNA oligonucleotide (IDT) complementary to the 3’ constant region of the libraries, to prevent participation of the constant region in the binding reaction (52), as is commonly done for ribozyme selections (53).

### Single-target equilibrium binding assays

Target RNAs that had been radiolabeled (as described in supplementary methods) were diluted in Buffer R (18 mM HEPES pH 7.4, 100 mM potassium acetate, 1 mM magnesium acetate, 0.01% IGEPAL-630 (Sigma-Aldrich, I3021), 0.01 mg/mL yeast tRNA (Life Technologies, AM7119), 5 mM DTT (Invitrogen, 15508-013), and 1 U/μL SuperaseIn (Invitrogen, AM2694)) to 10/6 times the desired final concentration in the binding reactions (1–5 pM). A single-use aliquot of AGO–miR prep was thawed and dilutions were prepared in Buffer DB (18 mM HEPES pH 7.4, 100 mM potassium acetate, 1 mM magnesium acetate, 0.01% IGEPAL CA-630, 0.1 mg/mL BSA (New England Biolabs, B9000), 0.01 mg/mL yeast tRNA, 0.1 μg/μL 3X FLAG peptide, 5 mM DTT, 20% glycerol) to 10/4 times the desired concentration for each binding reaction.

Binding reactions were then initiated by adding 6 μL of the RNA in Buffer R to 4 μL of AGO–miR in Buffer DB. After incubating the binding reactions at 37°C for 4 h, AGO–miR-bound target RNA was separated from free target RNA by nitrocellulose filter binding. Nitrocellulose disks (Amersham Protran 45 um Nitrocellulose (Cytiva, GE10600002) and nylon disks [either Amersham Hybond-XL (Cytiva, discontinued), Hybond-NX (Cytiva, discontinued) or Hybond-N+ (Cytiva, RPN303B)] were cut from membrane sheets using a hole punch with a diameter of 0.5 cm and pre-incubated in Buffer FB (18 mM HEPES pH 7.4, 100 mM potassium acetate, 1 mM magnesium acetate) at 37°C for at least 15 min prior to filter binding. A membrane stack (Hybond on the bottom, nitrocellulose on top) was assembled on a pedestal (Whatman filter holder, Sigma-Aldrich, WHA420100) on a Visiprep vacuum manifold (Sigma-Aldrich, 57250-U). After applying vacuum and checking the membrane stack had sealed to the pedestal by adding 10 μL of Buffer FB to the stack and observing it flow through, the contents of the 10 μL binding reaction were pipetted onto the membrane stack and allowed to flow through for a few seconds, before washing with 100 μL of ice-cold Buffer FBW (1X Buffer FB supplemented with 5 mM DTT). Washed membrane stacks were removed from vacuum, air-dried, and separated, and radioactivity was quantified by phosphorimaging (Amersham Typhoon Phosphorimager). The fraction of AGO–miR-bound RNA in each binding reaction was quantified by measuring the background-normalized intensity of the signal from the RNA captured on the nitrocellulose filter, and dividing that by the total background-normalized intensity from both the RNA captured on the nitrocellulose filter and that captured on the Hybond filter. Fraction bound (*θ_B_*) was plotted against the concentration of AGO–miR in the binding reaction and fit to the fractional saturation equation (equation 1) to extract *K*_d_ values.

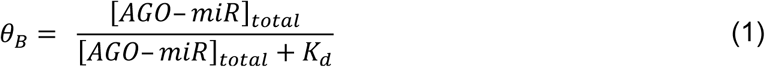

### Association rate assays

Equimolar mixtures of multiple 5′-^32^P-labelled target RNAs (radiolabeled as described in supplementary methods), each at 10/6 times their final reaction concentration of 10 pM, were prepared in Buffer R. As for the equilibrium binding assays, a dilution series of AGO–miR was prepared in Buffer DB, with each dilution at 10/4 times the desired final concentration of AGO– miR in the binding reaction. Binding reactions were initiated by mixing 60 μL of the target RNA mixture with 40 μL of the AGO–miR dilution. To maintain pseudo-first-order conditions, the lowest concentration of AGO–miR for which these binding reactions were conducted was always ≥80 pM. At each incubation time, a 10 μL aliquot of reaction mixture was subjected to filter binding. After air-drying of membranes and quantification of the total fraction of RNA bound at each timepoint, bound RNAs were eluted from the nitrocellulose membranes by Proteinase K (Invitrogen, 25530049) digestion in 420 µL Proteinase K Buffer (50 mM Tris-HCl, pH 7.4, 50 mM sodium chloride, 10 mM EDTA, 1% SDS) for 45 min at 60°C. The eluted RNA was then phenol-chloroform extracted (Phenol:chloroform:isoamyl alcohol (25:24:1), Sigma-Aldrich, P2069), ethanol precipitated, and re-suspended in 10 μL Gel Loading Buffer II (Invitrogen, AM8547). In parallel, some input RNA mixture was subjected to the same Proteinase K digestion, phenol-chloroform extraction, and ethanol precipitation. Input RNAs and eluted RNAs from each time point were resolved on a denaturing 15% polyacrylamide gel and quantified by phosphorimaging. The target RNAs (which differed from each other by ≥4 nucleotides) were well-resolved.

The background-normalized intensity of each band was plotted against time to track the appearance of each ternary AGO–miR–target complex as the reaction progressed. The signal for each band was further normalized to its input value and to the total fraction bound at that timepoint as quantified by phosphorimaging of the membranes before Proteinase K digestion. Because AGO–miR was in excess over the mixture of target RNAs (and therefore over each individual target), pseudo-first-order conditions were assumed, and the production of each AGO–miR–target complex over time was fit to the integrated rate equation for pseudo-first-order irreversible bimolecular association (equation 2), in which *AR*(*t*) is the molar concentration of AGO–miR–target complex at time *t*, *AR*_eq_ is the molar concentration of AGO–miR–target complex once the reaction has reached equilibrium, *A*_T_ is the total molar concentration of AGO– miR in the reaction, and *k*_on_ is the elemental rate constant for association (units of M^−1^s^−1^).

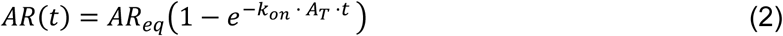

### Hydroxyl radical probing

Because AGO–miR preps produced for hydroxyl radical probing experiments featured radiolabeled guides, their concentration could not be easily quantified by measuring binding to a radiolabeled 7mer-m8 target as described in supplementary methods. Thus, quantification was performed by running an aliquot of AGO–miR prep on a denaturing 20% polyacrylamide gel, alongside standards with known amounts of radiolabeled guide. 6 μL aliquots of AGO–miR at 1– 3 nM were prepared in Buffer S– (18 mM HEPES pH 7.4, 100 mM potassium acetate, 1 mM magnesium acetate, 0.01% IGEPAL-630, 0.01 mg/mL yeast tRNA, 5 mM DTT, and 0.1 mg/mL BSA), and 9 μL aliquots of target RNAs at 50 nM were prepared in Buffer R– (Buffer R without glycerol). AGO–miR and target RNA (or buffer-only control) aliquots were mixed and pre-incubated at 37°C for 30 min before hydroxyl radical probing. To initiate probing, 2 μL of Mix F (13.33 mM ammonium ferrous (II) sulfate (Sigma-Aldrich, 09719), 14.667 mM EDTA pH 8, 100 mM DTT) was added to the 15 μL binding reaction mixture, followed by 3 μL of Mix X (1% hydrogen peroxide (Hydrogen Peroxide (30%), Sigma-Aldrich, 216763) in 2.5 X Buffer RF (Buffer R with 9mM magnesium acetate instead of 5mM magnesium acetate)). After 2 min at 37°C, probing reaction mixtures were quenched by the addition of 5 μL of ice-cold 500 mM sodium sulfite (Sigma-Aldrich, S0505). After at least 5 min of quenching on ice, samples were phenol:chloroform extracted, and RNA was ethanol precipitated. Cleavage products were resolved on a denaturing 20% polyacrylamide sequencing gel. Negative-control pre-quenched samples were also included to establish a baseline for non-specific cleavage of the guide RNA during post-probing workup. For pre-quenching, mixes X and F were combined, reacted for 2 min, and quenched for 5 min before being added to the binding reaction. Positive-control samples of free guide were used to establish a ceiling for the degree to which each position along the guide could be cleaved when it was maximally solvent accessible. For probing free guide, aliquots of AGO–miR were extracted with phenol:chloroform, ethanol precipitated (with no co-precipitants), resuspended in Buffer S– and subjected to hydroxyl radical probing. For each band, the fraction of signal in the lane attributed to the band was measured, and relative·OH reactivity at each position was calculated by the formula (*S* – *Q*)/(*N* – *Q*), in which *S* is that fraction of total signal attributed to the band, and *Q* and *N* are the equivalent values for that cleavage product in the pre-quenched negative-control and free-guide positive-control, respectively. Relative ·OH reactivity could not be reliably calculated for all positions of all guides, as discussed in the supplementary methods.

### Periodate probing of 3’-end accessibility

Periodate probing of guide 3′-end accessibility in AGO–miRNA complexes was conducted according to a protocol modified from those used for complete conversion of 3′-ends (54), with the goal of achieving partial reactivity and compatibility with protein−RNA complexes. 1 nM AGO–miR (or free guide that had been extracted with phenol:chloroform, as was done for hydroxyl radical probing experiments) in Buffer S– was incubated with 4 nM target RNA (or water) in a borate-supplemented reaction buffer (22.5 mM HEPES pH 7.4, 62.5 mM borate pH 8.9, 125 mM potassium acetate, 1.25 magnesium acetate, 6.25 mM DTT, 0.025 mg/mL BSA, 0.0125 mg/mL yeast tRNA, and 0.0125% IGEPAL CA-630) that provided a final pH of 8.5. After 30 min pre-incubation at 37°C, 25% volume of pre-warmed sodium periodate (Sigma-Aldrich, 71859-25G) was added (to a final concentration of 10 mM) to initiate probing, which was allowed to proceed at 37°C for 20 seconds. The reaction was then quenched for at least 5 min at 4°C with 100% volume of ice-cold quenching solution (100 mM sodium sulfite, 200 mM Tris pH 7.5). For pre-quenched negative controls, the periodate solution was mixed with the quenching solution for 5 min before adding AGO–miR. 20 µL of each quenched reaction was mixed with 380 µL of cold 0.3 M sodium chloride and 5 µL of 15mg/mL GlycoBlue co-precipitant (Invitrogen, AM9515), and precipitated with 1 mL of ethanol. Dried pellets were resuspended in 10 µL of β-elimination buffer (100 mM borate pH 9.5) by incubation at 37°C for 15 minutes, then incubated at 45°C for 90 minutes. 10 µL of 200 mM Tris-HCl and 25 nM non-radiolabeled guide in 0.75X Gel Loading Buffer II was added to each sample, and samples were resolved on a denaturing 20% polyacrylamide sequencing gel, which was then phosphorimaged. Relative periodate reactivity was quantified as described above for relative ·OH reactivity in hydroxyl radical probing experiments.

### Massively parallel reporter assays (MPRA)

Guide and passenger strand RNA oligos (IDT) were annealed to make miRNA duplexes for miR-155, miR-124, and miR-1. For miR-155, 10 μM duplex was annealed in Low-Salt Annealing Buffer (20 mM Tris-HCl pH 7.5, 40 mM sodium chloride, 2 mM EDTA) to avoid G-quadruplex formation, but for the other two miRNAs, annealing was conducted in Annealing Buffer (60 mM Tris-HCl pH 7.5, 200 mM sodium chloride, 2 mM EDTA). Annealing reactions were incubated for 4–5 min at 100°C on a benchtop metal heat block, then slowly cooled by removing the block from heat and allowing it to cool to ∼30°C, at which point, tubes were transferred to ice for 1 h, then stored at –20°C. Annealing was confirmed by analysis on native polyacrylamide gels, run in a cold room (5.5°C) and stained with SYBR gold (Invitrogen, S11494).

HeLa or F9 cells were grown in 10-cm plates to ∼50% confluence before being co-transfected with plasmid and miRNA duplex using Lipofectamine 2000 (Invitrogen, 11668019) according to manufacturer’s instructions. Each transfection had 5.8 μg of plasmid library (for single-site experiments, 5.8 μg of single-site library; for dual-site experiments, 2.9 μg of single-site library and 2.9 μg of dual-site library) along with 28.9 μg of carrier DNA (pUC19, New England Biolabs, N3041) and miRNA duplex, such that the final concentration of miRNA duplex in the media was 25 nM. Plasmid libraries were prepared as described in supplementary methods. For mock-transfected samples, the procedure was the same, except an identical volume of annealing buffer was added instead of miRNA duplex. After 24 h, media was aspirated, and cells were washed twice with 5 mL ice-cold PBS before being lysed by resuspension in 362 μL of lysis buffer (10 mM Tris-hydrochloride pH 7.5, 5 mM magnesium chloride, 100 mM potassium chloride, 1% v/v Triton X-100 (Invitrogen, HFH10), 2 mM DTT, 0.02 U/μL SuperaseIn, 1 tab / 10 mL cOmplete mini EDTA-free protease inhibitor cocktail (Roche, 11836170001)) and gentle shearing by passing four times through a 26 G needle. Debris was removed by centrifugation for 10 min at 1300*g* at 4°C, and the supernatant from each plate was split into three 150 μL aliquots for RNA extraction with Trizol-LS (Invitrogen, 10296010) according to the manufacturer’s instructions, after which the three samples from each plate of cells were pooled. For each plate, 10 μg of extracted RNA sample was treated with TURBO DNase (Invitrogen, AM2238) according to manufacturer’s instructions, phenol:chloroform extracted, ethanol precipitated, and resuspended in 10 µL water. 5 μL of each sample was then reverse transcribed using Superscript III Reverse Transcriptase (Invitrogen, 18080093) and an RT primer specific to the 3′ constant region of the reporter mRNA, in a 32 µL reaction, after which RNA was hydrolyzed using 1 M sodium hydroxide at 90°C for 10 min before neutralization with 1M HEPES pH 7.4. 8 μL of each cDNA was amplified using Phusion polymerase (New England Biolabs, M0530) and barcoded sets of primers containing Illumina P5 and P7 sequences that were complementary to the 5’ and 3’ constant regions of the MPRA libraries, to generate libraries for sequencing on an Illumina NovaSeq SP flow cell. Amplified libraries were size-selected on 2.5% Metaphor agarose gels (Lonza, 50181) and extracted (QIAquick Gel Extraction Kit, Qiagen, 28704). Each replicate of an MPRA experiment began with four plates of cells, respectively co-transfected with either miR-155, miR-124, or miR-1 (a negative control), or mock-transfected with annealing buffer in place of a miRNA duplex. The second replicate for an experiment was started a few days later with another four plates of the same cell line grown from the same early-passage frozen stock.

### Analysis of AGO-RBNS data

Reads were filtered to remove those with uncertain base calls and those for which the last four nucleotides did not match the 3′ constant region of AGO-RBNS libraries. For each sample, reads were assigned to individual miRNA sites as described in supplementary methods. Reads that did not contain any sites to the relevant miRNA were assigned to the no-site background. Tables of numbers of reads assigned to each site and the no-site background across the different concentrations of AGO–miR and in input libraries were used to fit relative *K*_d_ values as described (28). Relative *K*_d_ values describe how much better a site was bound compared to the no-site background. Negative log-transformed relative *K*_d_ values for the many compound sites that featured both 3′ complementarity and seed complementarity were fit to a multiplicative model (equation 3) wherein independent 3′ complementarity, seed complementarity, and offset coefficients summarized the typical contribution to binding of individual patterns of 3′ complementarity, seed complementarity, and offset, across the many different combinations in which they appear.

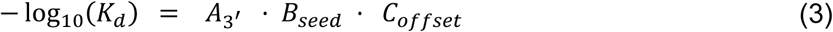

### Analysis of MPRA data

Reads from single-end sequencing of final libraries from HeLa cell single-site MPRA experiments were filtered to just those reads with no uncertain base calls, then quality-filtered. Reads were assigned to library variants if the entire sequence of the variable region was a perfect match to one of the library variants. Counts per million (CPM) values were calculated for each site across the multiple contexts in which it appeared, and fold-change values for each site were calculated for the miRNA-transfected versus mock-transfected samples, normalized to the fold-change values observed for no-site variants. Reads from paired-end sequencing of final libraries from F9 cell single-site MPRA experiments were joined before being identically filtered, assigned, and quantified. For the F9 cell dual-site MPRA experiment, despite paired-end sequencing, read 1 alone was used to assign reads if the first 150 nucleotides of the read matched the first 150 nucleotides of the variable region of one of the library variants.

### Survey of endogenous miRNA target sites

Representative isoforms for human 3′ UTR sequences from TargetScan 7.2 (31) were searched for sites to the guide strands of the 183 confidently annotated human miRNAs from the 108 seed families that are conserved to zebrafish according to TargetScan 8 (28), using the same logic employed to assign AGO-RBNS reads to different miRNA sites described in the supplementary methods. Site identities and positions within 3′ UTRs were recorded for all miRNAs and all UTRs, even for sites for which it was difficult to unambiguously assign site identity. Canonical sites with 6mer, 7mer-A1, 7mer-m8 or 8mer seed matches (with or without supplemental 3′ complementarity) were counted, as were 3′-only sites with at least 10 nucleotides of contiguous 3′ complementarity and less than three nucleotides of contiguous seed matching, and 3′+microseed sites with at least 10 nucleotides of contiguous 3′ complementarity and 3–5 contiguous nucleotides of seed matching at an offset of between –4 and +6.

### Analysis of mRNA-seq data from miRNA transfections

Analysis was on RNA-seq data measuring the response of endogenous mRNAs following transfection of one of 16 of miRNAs into HeLa cells (28). Reads were filtered to remove any with uncertain base calls before aligning to hg19 using STAR, providing splice junction and transcript annotations (55). Read counts per gene were determined using featureCounts (56), and converted to batch-normalized log_10_-transformed transcripts per million (logTPM) values as described (28). Log_2_(fold-change) values for each gene were calculated for each miRNA relative to the mean of its logTPM value in the other 15 miRNA-transfected samples. For each miRNA, cohorts of genes whose dominant 3′ UTR isoform (see supplementary methods) contained a single 7mer-m8 site or a single ≥8-nt 3′-only site were assembled, along with a cohort of UTR length-matched no-site controls (see supplementary methods). Cumulative distribution functions (CDF) for the log_2_(fold-change) values for those cohorts for each miRNA were constructed. CDFs for 7mer-m8 and 3′-only cohorts were compared to CDFs for no-site control cohorts by the two-sided Kolmogorov–Smirnov (KS) test, to determine whether single 7mer-m8 and 3′-only sites imparted changes to their respective mRNAs.

## DATA AVAILABILITY

Sequencing data from AGO-RBNS and MPRA experiments have been deposited at National Center for Biotechnology Information Gene Expression Omnibus (accession number GSE290589).

## CODE AVAILABILITY

New code generated for this study is deposited at Zenodo (doi: 10.5281/zenodo.14766495). For additional relevant resources and help with implementation, please contact first author Matthew Hall.

## SUPPLEMENTARY MATERIALS

Supplementary materials are available online

## AUTHOR CONTRIBUTIONS

M.H.H and D.P.B conceived the project, designed the study, and wrote the manuscript. M.H.H performed experiments and associated data analysis. P.Y.W and T.M.P contributed to project conceptualization, interpretation of results, editing of the manuscript, and contributed new code for use in data analysis. P.Y.W conducted and analyzed periodate probing experiments, and conducted miR-124 hydroxyl radical probing experiments.

## ACKNOWLEDGEMENTS

We thank S.E. McGeary for original AGO-RBNS analysis code. We thank S.E. McGeary, K. Xiang, R. Muller and others in the Bartel laboratory for helpful discussions. We thank the Whitehead Genome Technology core for help with sequencing experiments.

