## Supplementary Information for "Functional microRNA targeting without seed pairing"

### SUPPLEMENTARY METHODS

#### Preparation of radiolabeled RNAs

Guide and target RNAs to be radiolabeled were synthesized (IDT) with either a 5'-OH (target RNAs) or a 5'-monophosphate (guide RNAs), gel-purified on a denaturing 15% polyacrylamide gel, ethanol precipitated, and resuspended in water. 5'-phosphorylated guide RNAs were dephosphorylated using shrimp alkaline phosphatase (rSAP) (New England Biolabs, M0371) at 37°C for 30 min, followed by heat inactivation at 75°C for 5 min, desalting on Micro Bio-Spin P-6 columns (Bio-Rad 7326221), and a second ethanol precipitation. Radiolabeling was performed by incubating 1–10 pmol guide or target oligonucleotide with 0.33 U/µL T4 Polynucleotide Kinase (New England Biolabs, M0201) and 1–3 µL per 10–12 µL reaction of [ɣ-$^{32}$P] ATP (Revvity, BLU002Z250UC) for 1.5 h at 37°C. After radiolabeling of guide RNAs, a 15 min chase with 0.16 mM cold ATP (Thermo Scientific, R0441) was conducted. Reaction mixtures were column-purified on Micro Bio-Spin P-6 columns, gel-purified on a denaturing 15–20% polyacrylamide gel, ethanol precipitated, and re-suspended in water to the desired stock concentration (1–10 nM), assuming a 50% loss of product during gel purification.

#### Quantification of purified AGO–miR preps

At time-of-use, the concentration of functional AGO–miR in thawed single-use aliquots of AGO–miR prep was quantified by a binding assay, using a target RNA that contained a single 7mer-m8 site to the guide RNA. Target RNA (1.6 nM) in Buffer R (18 mM HEPES pH 7.4, 100 mM potassium acetate, 1 mM magnesium acetate, 0.01% IGEPAL-630, 0.01 mg/mL yeast tRNA, 5 mM DTT, and 1 U/µL SuperaseIn) was prepared as a 1:10 mixture of 5′-$^{32}$P-labelled and unlabeled RNA. Serial dilutions of the aliquot of AGO–miR were prepared in Buffer S (18 mM HEPES pH 7.4, 100 mM potassium acetate, 1 mM magnesium acetate, 0.01% IGEPAL-630, 0.01 mg/mL yeast tRNA, 5 mM DTT, 0.1 mg/mL BSA, and 25% glycerol), and 4 µL of each added to 6 µL of target RNA to give a final concentration of target RNA in the binding reactions of 1 nM. Binding reactions were incubated at 37°C for 2–4 h, before filter binding. As expected, a linear increase in fraction bound was observed at low AGO–miR concentrations. Linear fitting to the 3–5 datapoints at low concentrations enabled determination of the concentration of active AGO–miR.

#### Discussion of the inability to quantify relative •OH reactivity for all guide positions in hydroxyl radical probing experiments

There were two reasons why relative •OH reactivity scores could not be determined for all guide positions for all hydroxyl radical probing experiments. Firstly, trimming of the miRNA tail in the AGO–miR preps (which was prevalent for miR-124 and less severe but still detectable for miR-155 and let-7) meant that relative •OH reactivity could not be quantified for the last few nucleotides of the guide. If there were bands corresponding to trimming intermediates and those bands were sufficiently intense, then the difference in signal for that band due to additional production of equal-length products by ·OH-mediated cleavage could not be distinguished from noise. Second, relative •OH reactivity at guide positions 1–3 could not be resolved because those 1–3-nt cleavage products ran with the salt front on the gel and were not resolved. In addition, for miR-155, relative •OH reactivity could not be determined for positions 4–7 as it was for positions 8–18. In the pre-quenched (Q) negative control samples where the abundance of those cleavage products was lowest, there were no visible bands at the expected position on the gel for those species. This lack of signal resulted from the fact that miR-155 consistently radiolabeled less effectively than miR-124 and let-7a. For other guide positions, relative •OH reactivity was quantified as described in Materials and Methods. However, for positions 4–7 of miR-155, background-normalized *Q* values could not be determined because there was no signal in excess of background. For these positions we therefore assumed *Q* values of zero, reducing the formula for calculation of relative •OH reactivity from *(S-Q)/(N-Q)* to the simple ratio *(S/N)*.

**Preparation of MPRA libraries**
Each MPRA library, synthesized as a SurePrint HiFi OLS DNA pool (Agilent, G7281A), was amplified (KAPA HiFi HotStart ReadyMix, Roche, 7958935001) with primers that added BstXI and BsrGI restriction sites. PCR product of desired length was purified on an agarose gel (QIAquick Gel Extraction Kit, Qiagen, 28704). 100 ng of purified PCR product was then double-digested with BstXI (New England Biolabs, R0113) and BsrGI-HF (New England Biolabs, R3575), and the restriction digest product was purified by Qiagen QIAquick PCR clean-up (QIAquick PCR Purification Kit, Qiagen, 28104). The mEGFP vector (1, 2) was similarly digested. Ligation reactions (T4 DNA ligase, New England Biolabs, M0202) were carried out with 150 fmol of library digest product and 50 fmol of backbone digest product in a 50 µL reaction incubated at 14°C, 16°C, 22°C and 25°C for 30 min at each temperature. The ligation reaction mixture was then ethanol precipitated without any co-precipitants, and 5 µL, containing 50% of the ligation reaction mixture was electroporated (Gene Pulser XCell Total, Bio-Rad, 1652660) into 100 µL of MegaX electrocompetent cells (Invitrogen, C640003) with the electroporator set to 2 kV, 200 Ω, 25 uF on Exponential Mode. Because electroporation cuvettes had a volume of ~20 µL, the 105 µL were split into five equal aliquots, which were each electroporated separately. Each aliquot was added to 200 µL of 37°C S.O.C. medium for recovery. For each library, the 1.1 mL of electroporated cells in recovery medium was collected, and each of the 5 cuvettes were washed with another 200 µL of S.O.C. to make a 2.1 mL culture, which was incubated with shaking at 225 rpm for 1 h at 37°C before plating onto one hundred 10-cm ampicillin agar plates. After incubation overnight at 37°C, the plates were scraped to collect the bacteria, which were concentrated by centrifugation at 3500 rpm for 15 min at 4°C, and the plasmid library was purified from this pellet (PureLink HiPure Expi Plasmid Megaprep Kit, Invitrogen, K210008XP).

**Analysis of AGO-RBNS data**
Pre-processing of sequencing reads. Random libraries featured a 48-nt region of variable sequence between 5′ and 3′ constant regions. Programmed libraries featured a 49-nt region of variable sequence (including the eight nucleotides of programmed 3′ complementarity) between the constant regions. Sequencing was on an Illumina HiSeq 2500 in single-lane Rapid Mode, with 53 cycles, resulting in 53-nt sequencing reads. Raw reads were filtered using cutadapt (3) to retain those that met the two criteria of i) ending with at least four nucleotides corresponding to the beginning of the 3′ constant region and ii) not containing any uncertain (N) base calls. In addition, quality filtering using fastq_quality_filter (from FASTX toolkit http://hannonlab.cshl.edu/fastx_toolkit/) was then used to remove any reads that contained bases with a quality score of B or lower (Phred-64 encoding).

Assigning reads to different miRNA target sites. First, each line was searched for seed matches (6mer-A1, 6mer-m8, 6mer, 7mer-A1, 7mer-m8, and 8mer sites). A hit for a given site required matching not only the FASTA sequence of the site itself, but also any conditions on the nucleotides around that segment of complementarity to the miRNA. For example, for miR-155, the sequence of a full 8mer seed match was AGCAUUAA, and thus a 6mer match had to conform to the pattern BGCAUUAB, where B indicates a C, G, or U, which prevented double counting of the 8mer site as both an 8mer and a 6mer site.

For each canonical site found in the line, a 20-nt (or less if the canonical site match was <20 nt from the start of the read) window upstream of the match was queried for 3′ complementarity of at least four contiguous nucleotides, using the same FASTA logic described above. For example, an m13_18 3′-supplementary match had to match positions 13 through 18 of the miRNA but not any other position in the 3′-region from nucleotides 9 through the end of the miRNA). We note that this analysis has two limitations. Firstly, stretches of 3′ complementarity with internal bulges will be recorded as an instance of the first contiguous stretch of 3′ complementarity. Secondly, stretches of imperfect 3′ complementarity with internal mismatches will not be counted. If no 3′ complementarity of at least four nucleotides was found, the site was recorded as a seed-only canonical site. For each possible 3′-supplementary match found by this search, the offset was calculated. Because the impact of additional 3′ complementarity on seed-matched site affinity is minimal for very negative or very positive offsets (2), 3′-supplementary matches occurring with offsets less than –4 or greater than +10 nt were ignored.

Sometimes, multiple 3′-supplementary matches with acceptable offsets were found within the 20-nt window upstream of a canonical site, making unambiguous assignment of site identity difficult. In these cases, the longest 3′-supplementary match was selected. If multiple 3′-supplementary matches of equivalent length remained, they were further filtered to just those that occurred within a more stringent range of offsets (miR-155, –3 through +3 nt; miR-124, 0 through +5 nt), which are more favorable (2). In the very rare cases where this filtering did not select a single 3′-supplementary match, either because more than one, or none of the 3′-supplementary matches of maximal length occurred with favorable offset, the site was flagged and excluded from downstream analyses. If only one 3′-supplementary match remained, then the identity of the site was recorded in the form “3' complementarity | offset | seed match” (e.g. m15_18|+1|m2_8). The position of the site within the read (start and end position) was also recorded.

This seed-based search found all sites with at least six contiguous nucleotides of seed complementarity. Following the search for seed matches, a second search of the read was conducted, searching for matches to the miRNA 3′ region that included at least the eight nucleotides of programmed 3′ complementarity (m13_20 for miR-155, m11_18 for miR-124). The logic of this search resembled that of the seed-based search. For each 3′-site match, a 20-nt window downstream (or <20-nt if the 3′-site match was less than 20 nucleotides from the end of the read) was queried for seed complementarity of at least two contiguous nucleotides, and offset was calculated for each possible partial seed match that had an offset to the 3′-site of between –4 and +10 nt. Some reads had multiple partial seed matches with acceptable offsets, and these were dealt with in the same way as multiple possible 3′-supplementary matches upstream of a canonical site, described above. The longest partial seed match was selected, and the list of partial seed matches filtered to include only those that occurred within the more favorable range of offsets. In the very rare cases in which multiple partial seed matches passed this filtering, unambiguous assignment of site identity was difficult, and the read was excluded from downstream analyses. Sites with ≥6-nt seed complementarity and ≥8-nt 3′ complementarity were identified by both the seed-based search and the 3′-based search, and thus these double-counts were collapsed to single counts.

This site-counting analysis yielded a table of the sites present within each read and their positions. If a read contained multiple sites, it could not be deduced whether the RNA molecule was bound because of the presence of site 1, site 2, or the combination of site 1 and site 2. Therefore, only reads that contained a single site were assigned to that site, resulting in a table of counts of how many reads contained a single example of each site. Reads that did not contain any sites were assigned to the no-site background.

Fitting of relative $K_d$ values for sites. Site-count tables were obtained for the samples corresponding to the different concentrations of AGO–miR in the initial AGO-RBNS binding reactions. From a concatenated table of site counts for each sample of bound RNA as well the input RNA and the estimated concentration of AGO–miR in each sample, the script Fit_Site_Kds_4.R (2) was used to infer a relative $K_d$ value for each site. For a detailed mathematical treatment of relative $K_d$ fitting, see (1). The relative $K_d$ value for each site indicates how much better that site bound AGO–miR compared to the no-site background.

Fitting compound site relative $K_d$ values to a multiplicative model of site affinity. The negative $\log_{10}$-transformed relative $K_d$ values for compound sites that bound better than background (relative $K_d$ < 0) were fit to a multiplicative model in which each $-\log_{10}(K_d)$ is assumed to be the product of a 3′-complementarity coefficient (e.g., $A_{m13\_22}$), a seed complementarity coefficient (e.g., $B_{m2\_8}$) and an offset coefficient (e.g., $C_{+2}$) (equation 1).

$$-\log_{10}(K_d) = A_{3'} \cdot B_{seed} \cdot C_{offset} \quad (1)$$

Each term represents the contribution of the component pattern of 3' complementarity, seed matching, and offset to the overall site affinity. These coefficients were fit globally for relative $K_d$ values of >5,000 compound sites and thus represent the typical contribution that each pattern of complementarity or offset makes to overall site affinity across the many site architectures in which it appears. Global fitting was conducted in Python using the GLM (generalized linear model) functionality of the package statsmodels (4). For each set of parameters (e.g., the 3' complementarity coefficients), fitted values were normalized such that the maximum value was 1.0 and the minimum value was 0, and 95% confidence intervals for the fit were scaled accordingly.

**Analysis of MPRA data**

For HeLa cell experiments using the single-site library, samples were sequenced with 200-nt single-end reads on a NovaSeq SP flow cell. Reads were quality filtered using Fastq_quality_filter (from FASTX toolkit http://hannonlab.cshl.edu/fastx_toolkit/) to retain those reads that both did not contain any uncertain (N) base calls and had a quality score of at least 20 across 90% of the sequenced nucleotides. For F9 cell experiments using the single-site library, samples were sequenced with 100x100-nt paired-end reads on a NovaSeq SP flow cell. Paired-end reads were joined using overlap of the sequenced portion of the two reads and Fastq-join (5), with a minimum perfect overlap of six nucleotides, before being identically filtered. For experiments using the dual-site library, samples were sequenced with 150x50-nt paired-end reads on a NovaSeq SP flow cell, and the 150 nucleotides of read 1 were used to assign reads to each variant. Reads from the dual-site experiment were quality filtered to retain those that did not contain any uncertain (N) base calls. Reads were then assigned to library variants, requiring that the entire sequence of the variable region (single-site experiments) or the first 150 nucleotides of the read (dual-site experiments) perfectly matched the sequence of one of the library variants and discarding all reads that were not a perfect match to a library variant. CPM values for each site were summed across the 20 sets of context sequences in which it appeared. Each miRNA-transfected sample was compared to the mock-transfected sample to calculate for each library member the final $\log_2$(fold-change) repression values ($R_s$), using equation 2, wherein $y_{s,+}$ is the summed CPM for site S in the miRNA-transfected sample, $y_{s,-}$ is the summed CPM for site S in the mock-transfected sample, $y_{ns,+}$ is the summed CPM for the five no-site library members in the miRNA-transfected sample, and $y_{ns,-}$ is the summed CPM for the five no-site library members in the mock-transfected sample).

$$R_s = \log_2\left(\frac{\left(\frac{y_{s,+}}{y_{ns,+}}\right)}{\left(\frac{y_{s,-}}{y_{ns,-}}\right)}\right) \quad (2)$$

Identification of slicing sites. For miR-124, compound sites with a combination of i) partial or full seed match, ii) +0-nt offset, and iii) either m9_19 or m9_22 3′ complementarity were suitable substrates for slicing catalyzed by AGO2–miR-124. To identify such slicing sites, we reasoned that for standard miRNA-mediated repression, +0-nt offset sites should be no more effective than positive-offset sites, whereas for sites that are efficiently sliced, +0-offset sites should be more effective than positive-offset sites. Therefore, if repression efficacy for the +0-nt offset version of the site was significantly better than that of the positive-offset version of the site ($p<0.05$, Tukey's post-hoc test, Bonferroni-corrected), the site was considered a slicing site. For experiments in F9 cells, for which we had four replicates, more sites passed this threshold for statistical significance than for experiments in HeLa cells, for which we had only two replicates.

Use of data from single-site variants in dual-site experiment as pseudo-replicates. First, the single-site MPRA experiment was conducted in F9 cells, and repression values determined for each of the 441 sites as described above, with 2 replicates. For the dual-site experiment, single-site and dual-site libraries were co-transfected into F9 cells, again with 2 replicates. Those 2 repression values for the single-site library variants in this experiment were considered as additional replicates for the initial 2 single-site repression values, bringing the total number of replicates for single-site repression values in F9 cells to 4.

**MPRA library design**

Flanking context sequences around site regions. For the single-site MPRA there was a 35-nt site region which was flanked by 50 nucleotides of context sequence on either side. 20 different pairs of upstream and downstream contexts were generated and appended each side of the site region, so that each of the 441 sites in the library was represented in 20 different sequence contexts, resulting in a total library complexity of 8,820 members. Each context sequence was generated with the following constraints. First, di-nucleotide frequencies had to match those determined for annotated HeLa 3′ UTRs (6). Second, seed complementarity of more than three contiguous nucleotides or 3′ complementarity of more than five contiguous nucleotides to either miR-155, miR-124, or miR-1 was not permitted. Third, 7mer-m8 matches to the 20 most highly expressed miRNAs in HeLa cells (7) were also prohibited, as were polyadenylation site (PAS) sequences (AWUAAA, where W indicates A or U), splice donor sequences (GGGURAGU, where R indicates A or G), BstXI and BsrGI recognition sequences ($CCAN_6UGG$ and UGUACA, respectively), sequences with potential to form G-quadruplexes, Pumilio sites (UGUANAUA), AU-rich elements (AUUUA), and homopolymeric stretches of more than five nucleotides. Experiments were first conducted in HeLa cells and then repeated in F9 cells, for which small RNA-seq data was not available to determine the 20 most highly expressed miRNAs. That said, site repression values from the two different cell lines correlated well (Fig. S7), which suggested that the different miRNA repertoire in F9 cells did not affect the ability of these library designs to report on repression by the transfected miRNAs.

Site region architectures. The central 35-nt site region of library members was split into a 15-nt 3'-site region upstream of a 20-nt downstream region that included an 8-nt seed-site region for canonical sites and compound sites. For 3'-only and compound sites, the 3'-site region consisted of the desired segment of 3' complementarity with the rest of the 15 nucleotides filled in with nucleotides that did not extend that 3' complementarity. For canonical sites and no-site variants, the 3'-site region did not have more than three nucleotides of contiguous pairing to miR-155, miR-124 or miR-1. For 3'-only sites, the 20-nt downstream region did not have more than one nucleotide of contiguous complementarity to the relevant seed. 20 of these downstream-region sequences that lacked seed complementarity were generated for each miRNA, so that each 3'-only site was represented in the context of 20 different downstream-region sequences. Matching the di-nucleotide composition of these downstream region sequences to that of HeLa 3′ UTRs, as was done for the 50-nt context sequences appended either side of the central 35-nt site region, was not possible due to the sequence constraints imposed by avoiding two contiguous nucleotides of seed matching.

For canonical sites and zero-offset compound sites, 20 downstream-region sequences were generated for each site in which the first eight nucleotides of that sequence had the desired seed matches, whereas the remaining 12 nucleotides did not extend the desired seed matches. For positive-offset compound sites the first two (miR-155 sites) or three (miR-124 sites) nucleotides of the downstream region were oligo(A), followed by the relevant 8-nt sequence that contained the desired seed match, and then 10 (miR-155) or 9 (miR-124) nucleotides of sequence that did not extend that desired seed match. As implemented for other generated sequences such as the 50-nt context sequences, and 20-nt flanking sequences appended downstream of 3′-only sites, 20 examples of these 12-, 10- or 9-nt flanking sequences were generated and appended for each seed match, so that each seed match was represented in 20 different sequence contexts. For each canonical site (6mer-A1, 6mer-m8, 6mer, 7mer-A1, 7mer-m8, 8mer), five different 35-nt site-region sequences were generated, each of which featured a different sequence in the 3'-site region, which did not contain more than two nucleotides of contiguous complementarity to either miR-155 or miR-124. For no-site variants, these same five no-site sequences per miRNA were used for the 3'-site region, and the 20-nt downstream region did not have more than two nucleotides of contiguous complementarity to either miR-155 or miR-124. For a detailed schematic of these MPRA library designs, see Fig S12A (single-site MPRA) and Fig S12B (dual-site MPRA).

<u>Iterative generation and checking of library variant sequences.</u> Sets of context and downstream-region sequences were iteratively generated until at least 30 sets had been identified wherein the sequences of the final library (in which all segments, including the 5' and 3' constant regions, had been concatenated) did not contain either ≥4 contiguous nucleotides of seed complementarity or ≥6 contiguous nucleotides of 3' complementarity to miR-155, miR-124, or miR-1 (apart from of the desired site), a PAS, a splice donor, a BstXI or BsrGI recognition site (apart from the two that were necessary for library cloning), or G-quadruplex forming sequences, each defined as described above. Candidate library sequences were then folded using RNAfold (8), and 20 sets of sequences that lacked long internal duplexes, particularly in the 35-nt site region, were chosen to be used for the final MPRA library. For the dual-site MPRA, the same design process was followed as for the single-site MPRA, with the same constraints. Each dual-site variant contained two identical 35-nt site regions, separated by an additional 20-nt spacer sequence generated with the same constraints as the 50-nt context sequences. The 35-nt site regions themselves were the same as those of the single-site MPRA.

**Survey of endogenous miRNA target sites**

The mature guide strand sequences of the 183 confidently annotated human miRNAs from the 108 seed families conserved to zebrafish were retrieved from TargetScan 8 (1). The hg19 genomic coordinates of all annotated 3′ UTR isoforms were retrieved from TargetScan7.2 (9), and the FASTA sequences corresponding to those coordinates were retrieved using bedtools (10). For each miRNA in the 183-miRNA cohort, each 3′ UTR FASTA sequence was searched for all examples of miRNA target sites with at least six nucleotides of contiguous seed complementarity, using the same logic described in the context of assigning reads to sites in AGO-RBNS analysis. This identified endogenous canonical sites (6mer, 7mer-A1, 7mer-m8, 8mer) and offset 6mer sites (6mer-A1, 6mer-m8), as well as any instances of upstream 3′-supplementary complementarity of at least four nucleotides that occurred within an acceptable offset range (–4 through +6 nt).

The 3′ UTRs were then searched for target sites with at least eight nucleotides of contiguous 3′ complementarity, using similar logic, and recording any instances of additional downstream seed complementarity of 3–5 contiguous nucleotides that occurred within the acceptable offset range of –4 through +6. When dealing with compound sites featuring both 3′ complementarity and seed complementarity, there were often multiple possible site identities, as when assigning sites to reads in AGO-RBNS analysis. The longest seed match was selected, and if there were multiple possible seed matches of the same length, the site was flagged and its identity left ambiguous but still counted as an example of a compound site. In addition to site identity, the start and end coordinates of the site within the 3′ UTR were recorded, as well as the identity of the 3′ UTR (in the form chr1:XXXX-YYYY, with XXXX and YYYY signifying the hg19 coordinates of the 3′ UTR, as retrieved from TargetScan 7.2).

For the purposes of assessing the relative abundance of different site types, canonical sites were not separated based on whether or not supplementary 3′ complementarity was observed upstream. Sites with ≥10 contiguous nucleotides of 3′ complementarity and <3 contiguous nucleotides of seed matching were classified as 3′-only sites, whereas sites with ≥10-nt contiguous 3′ complementarity and 3–5 contiguous nucleotides of seed matching were classified as 3′+microseed sites. As in AGO-RBNS analysis, compound sites with at least 6 nucleotides of seed complementarity were identified by both seed-based and 3′-based searches, and subsequently collapsed to single counts.

**Further discussion of mRNA-seq data analysis from miRNA transfections**

mRNA-seq data from HeLa cells transfected with a panel of 16 miRNAs (1) were retrieved in raw fastq format, quality filtered to retain reads without any uncertain (N) base calls, and aligned to hg19 using STAR, providing splice junction and transcript annotations (11) (options: --runThreadN 8, --outSAMtype BAM SortedByCoordinate –quantMode GeneCounts, otherwise all defaults). featureCounts (12) was used to count all single-mapping exonic reads (options: -d 15 -s 2 -T 12 -t exon -B -C) for each sample, and the tables of raw read counts were concatenated to generate a table for all samples. Read counts were converted to $\log_{10}$-transformed transcripts per million (TPM) values, and these logTPM values were batch-normalized by global fitting to a linear model (equation 3) wherein raw logTPM values were equal to a miRNA coefficient plus a batch coefficient. This fitting was accomplished using the ordinary least squares (OLS) function in the python module statsmodels (4). Because the replicates for a given miRNA could come from different batches, the contribution of the miRNA independent of batch effects is represented by $C(miR)$ (the miRNA-specific coefficient).

$$logTPM = C(miR) + C(batch) \tag{3}$$

The resultant table of batch-normalized logTPM values was then converted into a table of $\log_2$FC values for each miRNA relative to the mean of all the rest of the miRNAs (equation 4).

$$\log_2(m) = log_2\left(\frac{TPM_m}{\overline{TPM_n}}\right)\ for\ n \neq m \tag{4}$$

Of the 15,065 gene IDs for which reads were mapped by STAR and for which logTPM values were obtained, 12,181 had a single representative transcript ID associated with that gene ID in TargetScan 7.2 (9). For each of these 12,181 transcript IDs, coordinates of all 3′ UTR isoforms affiliated with that transcript ID were retrieved from TargetScan 7.2 (9). 10,511 had only a single 3′ UTR isoform affiliated with the transcript ID, and so the coordinates of those 3′ UTRs were retrieved. To conduct the previously described comprehensive survey of endogenous miRNA target sites, every 3′ UTR isoform annotated in TargetScan 7.2 was searched for sites to the 183 miRNAs from the 108 seed families conserved to zebrafish according to TargetScan 8 (1), and so the tables of sites to the 16 miRNAs of interest in those 10,511 3′ UTRs were retrieved.

For each of the 16 miRNAs, cohorts of transcripts from the 10,511 were assembled that contained a single 7mer-m8 site, a single 3′-only site, or no-site. First, the transcripts whose 3′ UTR contained only a single 7mer-m8 site and no other sites to the miRNA were selected as the positive-control cohort; for most miRNAs this included 100–250 transcripts. For each transcript in the positive-control cohort, 50 no-site transcripts belonging to the same 3′ UTR length bin (of 100 bins) were selected (with replacement) and added to the negative control no-site cohort. Then, the transcripts whose 3′ UTR contained only a single ≥10-nt 3′-only site and no other sites to the miRNA were selected as the experimental cohort. For each transcript in the experimental cohort, another 50 no-site transcripts from the same 3′ UTR length bin were selected (with replacement) and added to the negative control no-site cohort. Too few transcripts were present in these experimental cohorts for statistical analysis, and so transcripts whose 3′ UTR contained only a single ≥8-nt 3′-only site were selected, and the data from this analysis is depicted in Figure S11.

### SUPPLEMENTARY FIGURES

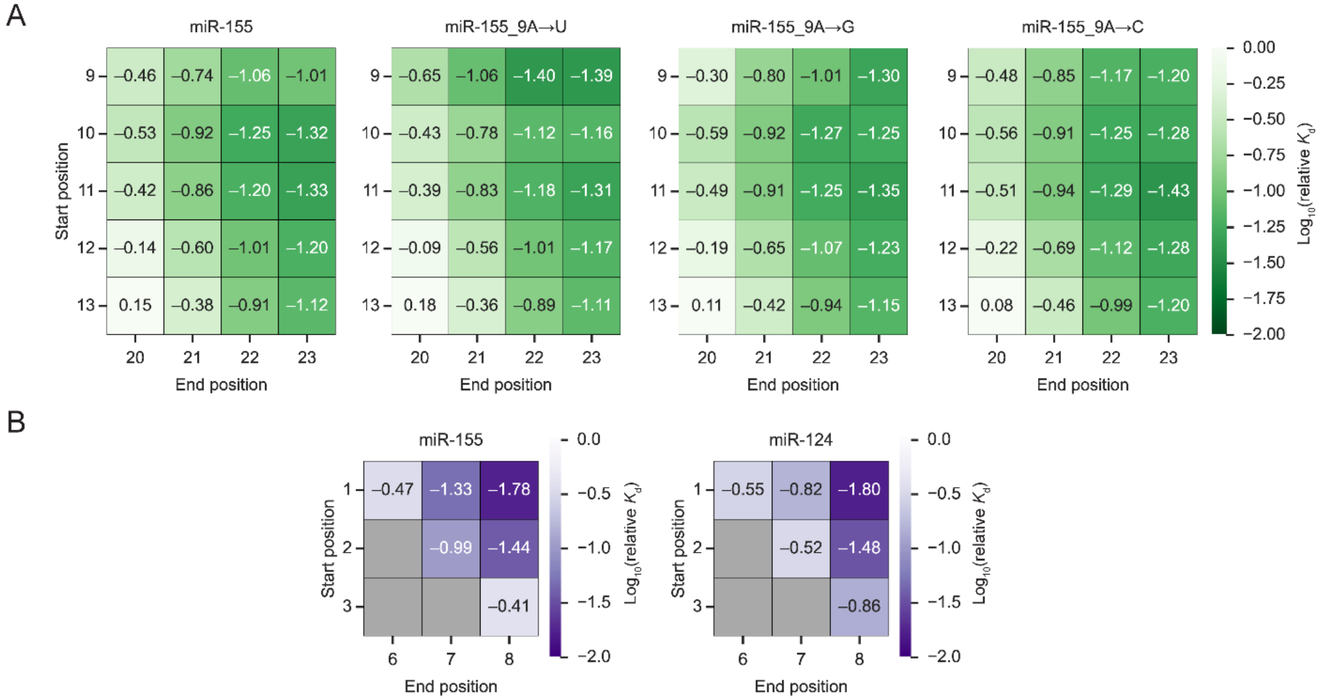


**Figure S1. miR-155 has preferences for the target nucleotide opposite position 9.** (A) The influence of the identity of the target nucleotide opposite position 9 of miR-155. Relative $K_d$ values were re-fit from the same AGO-RBNS data but specifying alternative miR-155 sequences that differed at position 9. Heatmaps depict $\log_{10}$-transformed relative $K_d$ values of different 3′-only sites. Panel 1 (far left) shows results for wild-type miR-155; panels 2, 3, and 4 show results for miR-155 with A at position 9 changed to U, G, or C, respectively. (B) Relative $K_d$ values of canonical sites. Heatmaps depict $\log_{10}$-transformed relative $K_d$ values of the indicated canonical sites of miR-155 (left) and miR-124 (right).

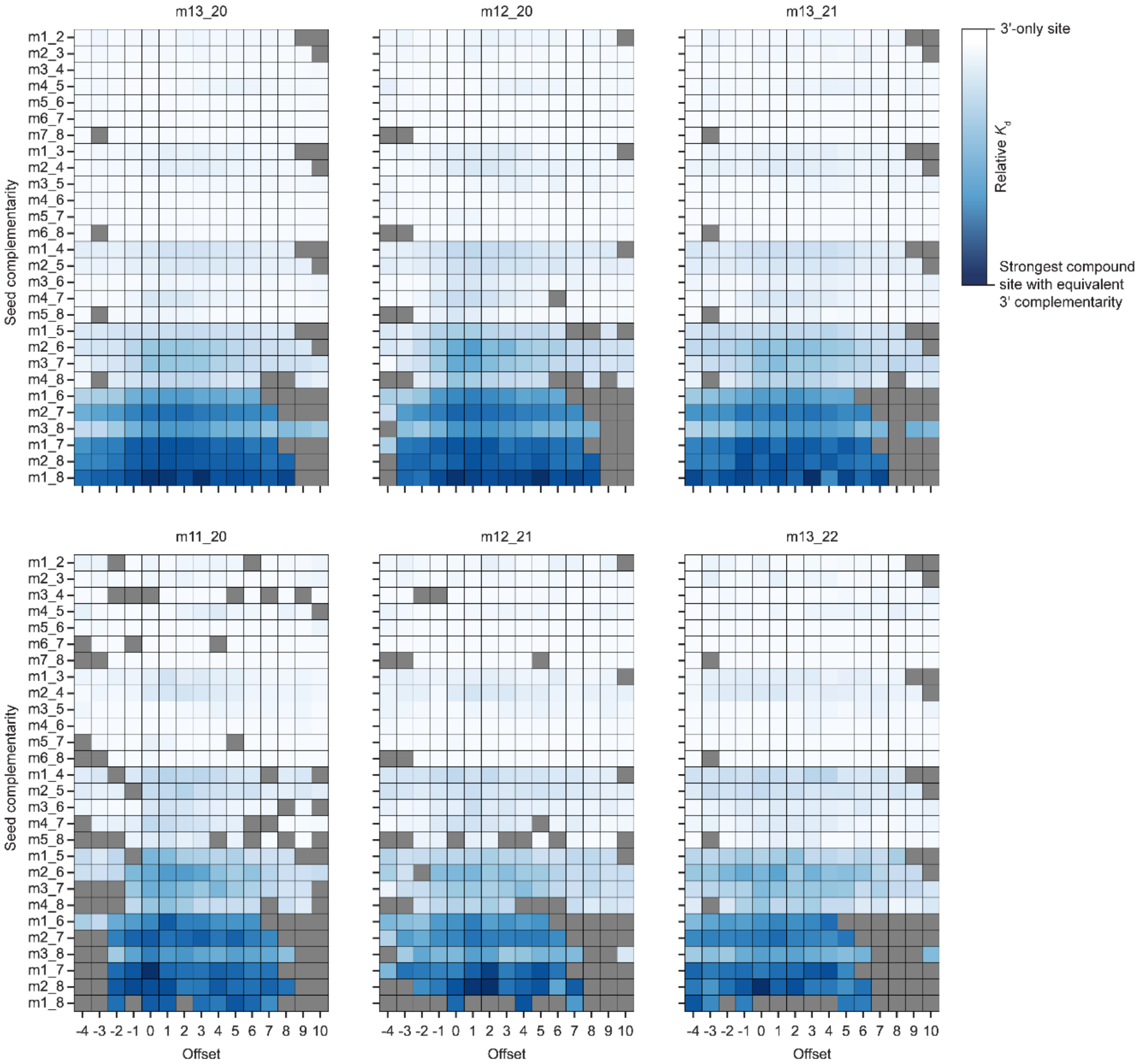


**Figure S2. Relative $K_d$ values for compound sites of miR-155.** Each panel depicts relative $K_d$ values for the indicated combinations of 3′ complementarity (panel title), seed complementarity (*y* axis), and offset (*x* axis). For each panel, the color is scaled such that the maximum relative $K_d$ value (white) indicates the relative $K_d$ of 3′ complementarity acting autonomously as a 3′-only site, and the minimum relative $K_d$ value (dark blue) indicates the highest-affinity relative $K_d$ attained across all combinations of seed complementarity and offset for that panel.

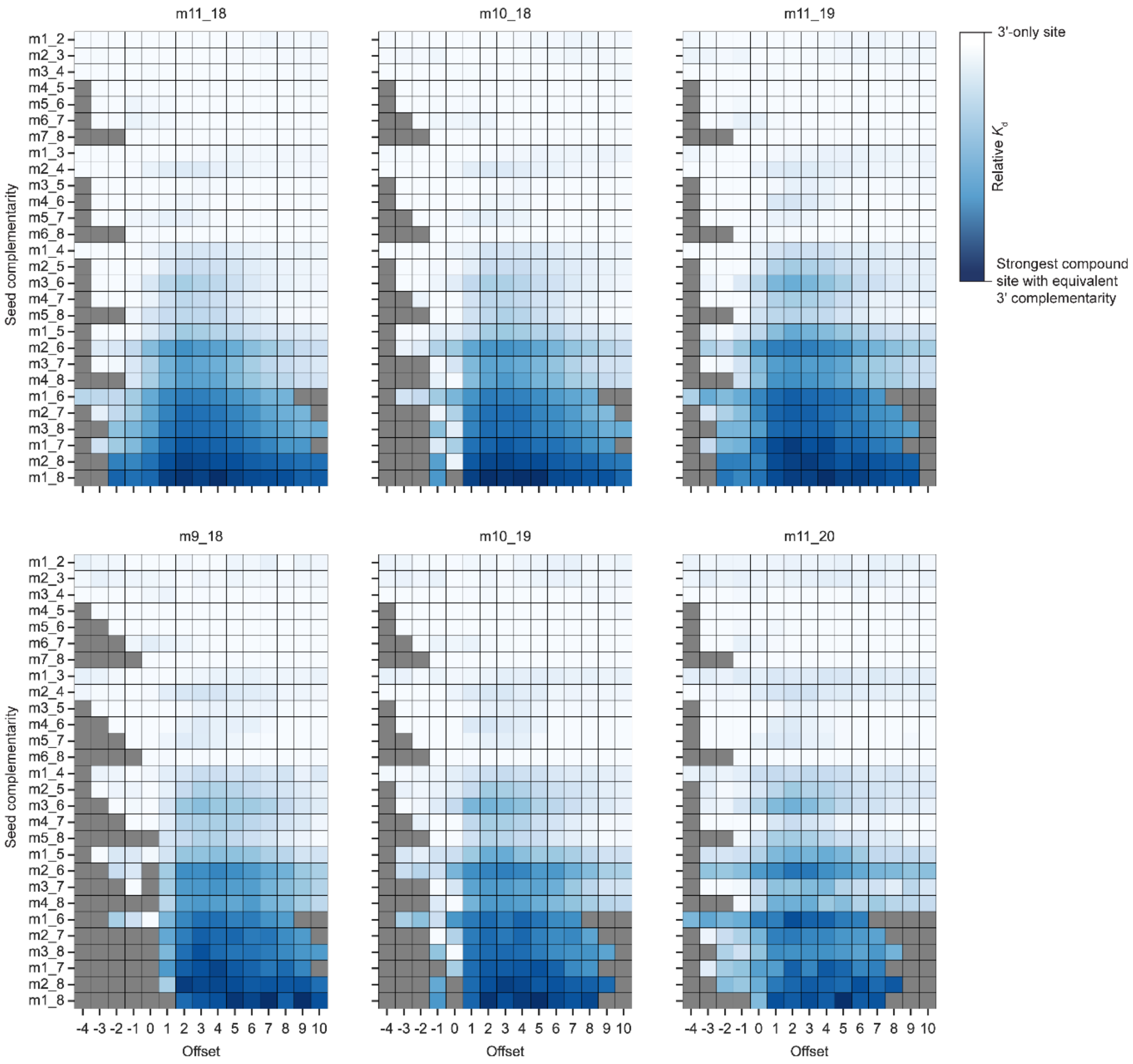


**Figure S3: Relative $K_d$ values for compound sites of miR-124.** Otherwise, this panel is as in Figure S2.

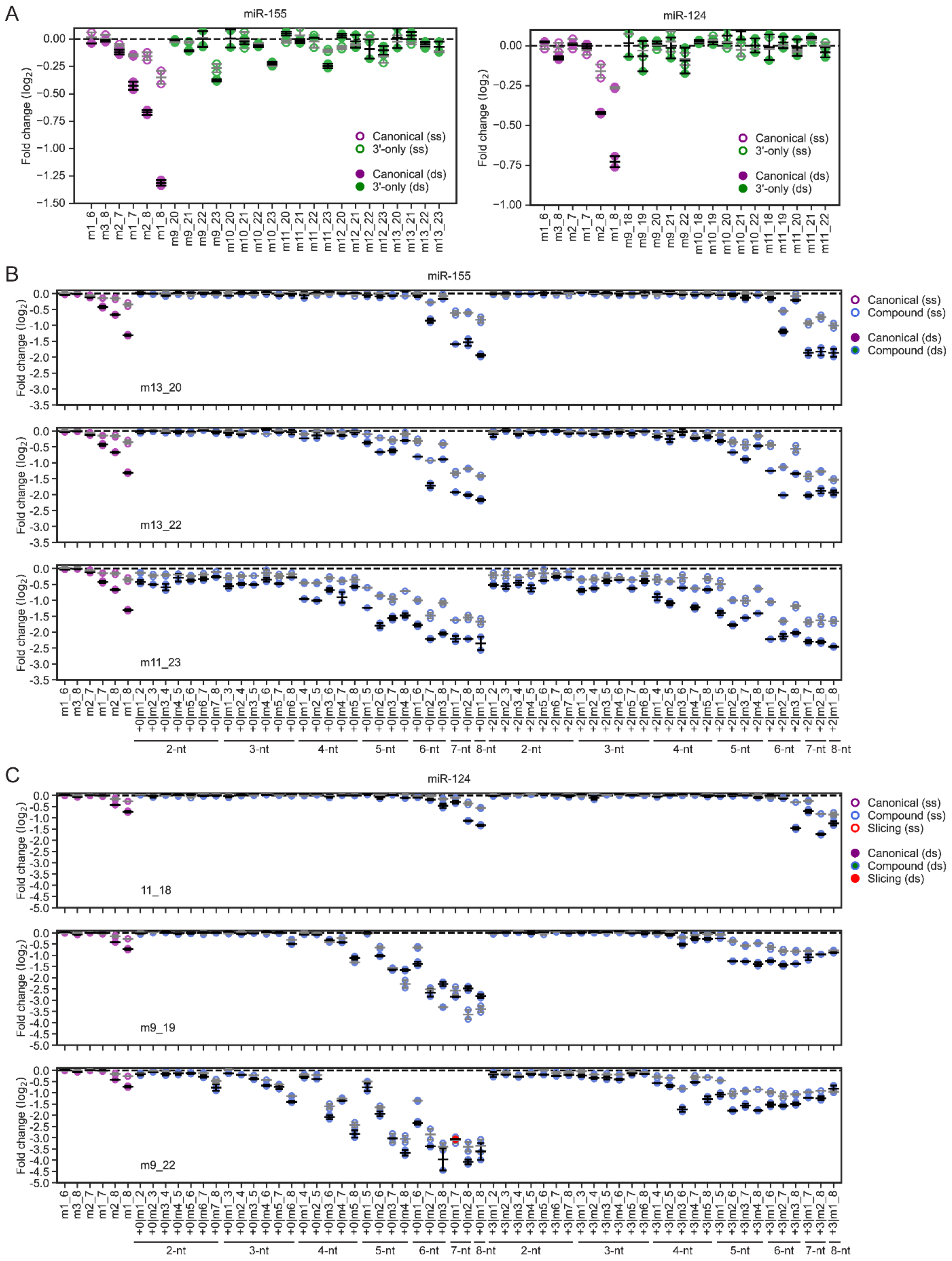
A
miR-155
miR-124
Fold change (log2)
Canonical (ss)
3'-only (ss)
Canonical (ds)
3'-only (ds)
B
miR-155
m13_20
m13_22
m11_23
Canonical (ss)
Compound (ss)
Canonical (ds)
Compound (ds)
2-nt
3-nt
4-nt
5-nt
6-nt
7-nt
8-nt
C
miR-124
11_18
m9_19
m9_22
Canonical (ss)
Compound (ss)
Slicing (ss)
Canonical (ds)
Compound (ds)
Slicing (ds)

**Figure S4. Closely spaced miRNA target sites generally act cooperatively.** (A) Repression efficacy of reporter RNAs with either single (unfilled) or dual (filled) sites of miR-155 (left) or miR-124 (right) upon transfection of the cognate miRNA. Otherwise, this panel is as in Figure 3C. *N* = 2 replicates per site (error bars, standard error of the mean). (B) Repression efficacy of reporter RNAs with either single (unfilled) or dual (filled) compound sites for miR-155 in response to miR-155 transfection. Otherwise, this panel is as in Figure 4A. *N* = 2 replicates per site (error bars, standard error of the mean). (C) Repression efficacy of reporter RNAs with either single (unfilled) or dual (filled) compound sites for miR-124 in response to miR-124 transfection. Otherwise, this panel is as in Figure 4B. *N* = 2 replicates per site (error bars, standard error of the mean).

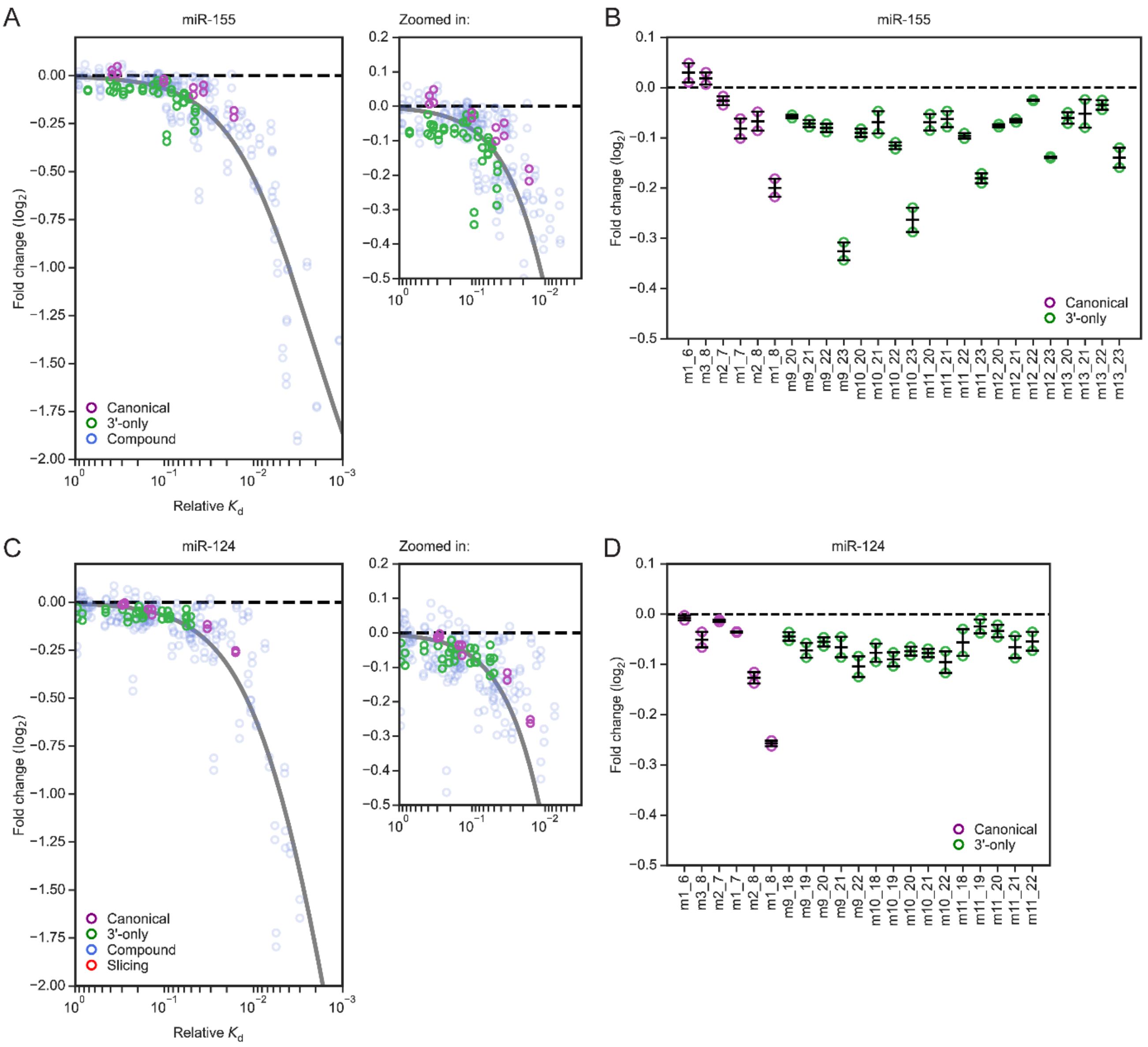


**Figure S5. 3′-only sites impart repression to reporter mRNAs in HeLa cells.** (A) Correspondence between repression of site-containing reporter mRNAs in response to miR-155 transfection in HeLa cells and the relative $K_d$ values of the sites measured by AGO-RBNS. $N$ = 2 replicates per site. Otherwise, this panel is as in Figure 3B. (B) Repression efficacy of canonical and 3′-only sites of miR-155 in HeLa Cells. $N$ = 2 replicates per site (error bars, standard error of the mean). Otherwise, this panel is as in Figure 3C. (C) Correspondence between repression of site-containing reporter mRNAs in response to miR-124 transfection in HeLa cells and the relative $K_d$ values of the sites measured by AGO-RBNS. $N$ = 2 replicates per site. Otherwise, this panel is as in Figure 3D. (D) Repression efficacy of canonical and 3′-only sites of miR-124 in HeLa cells. $N$ = 2 replicates per site (error bars, standard error of the mean). Otherwise, this panel is as in Figure 3E.

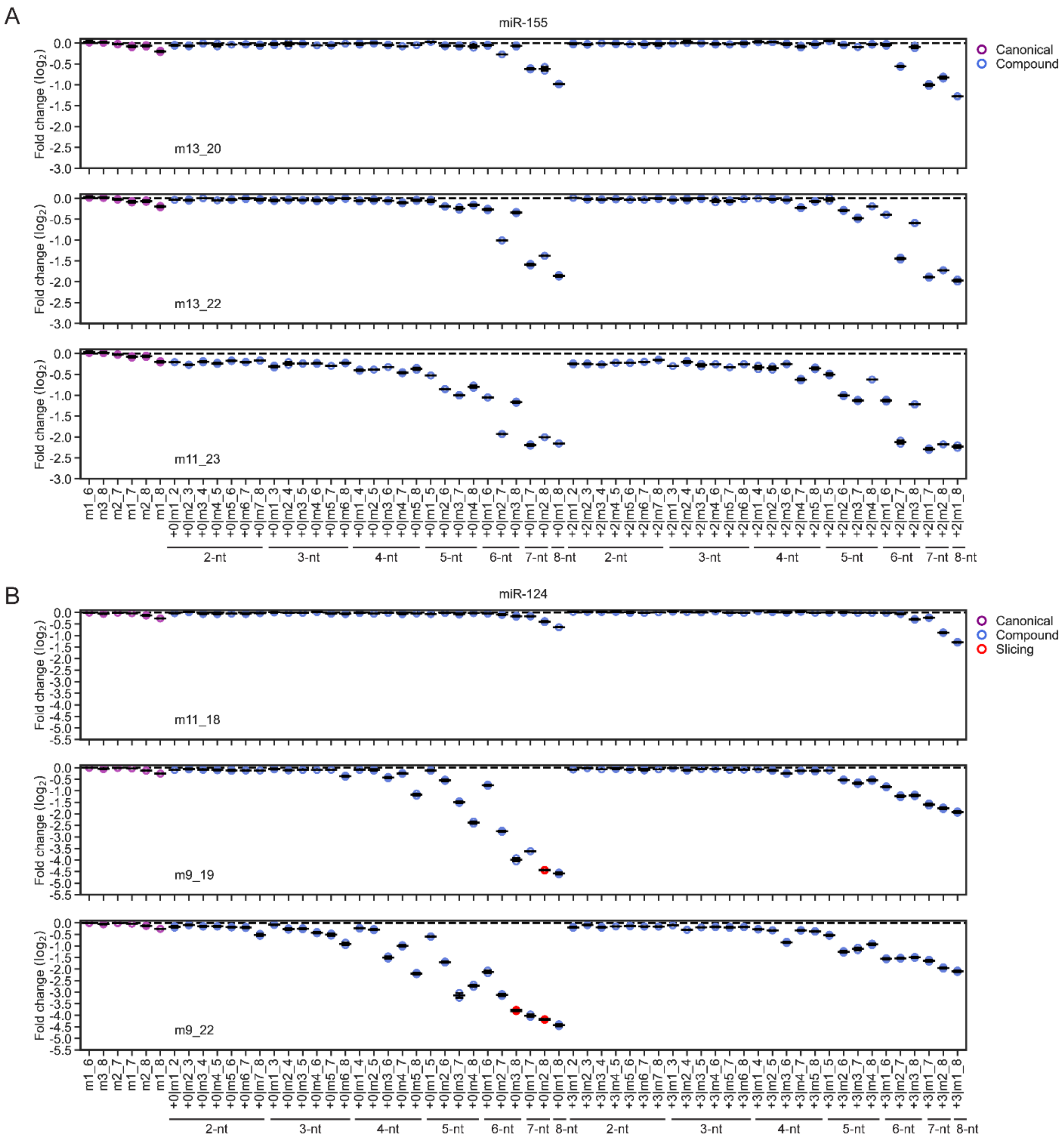


**Figure S6. Repression efficacy of compound sites in HeLa cells.** (A) Repression efficacy of reporter RNAs with compound sites of miR-155 in Hela cells. *N* = 2 replicates (error bars, standard error of the mean). Otherwise, this panel is as in Figure 4A. (B) Repression efficacy of reporter RNAs with compound sites of miR-124 in HeLa cells. *N* = 2 replicates (error bars, standard error of the mean). Otherwise, this panel is as in Figure 4B.

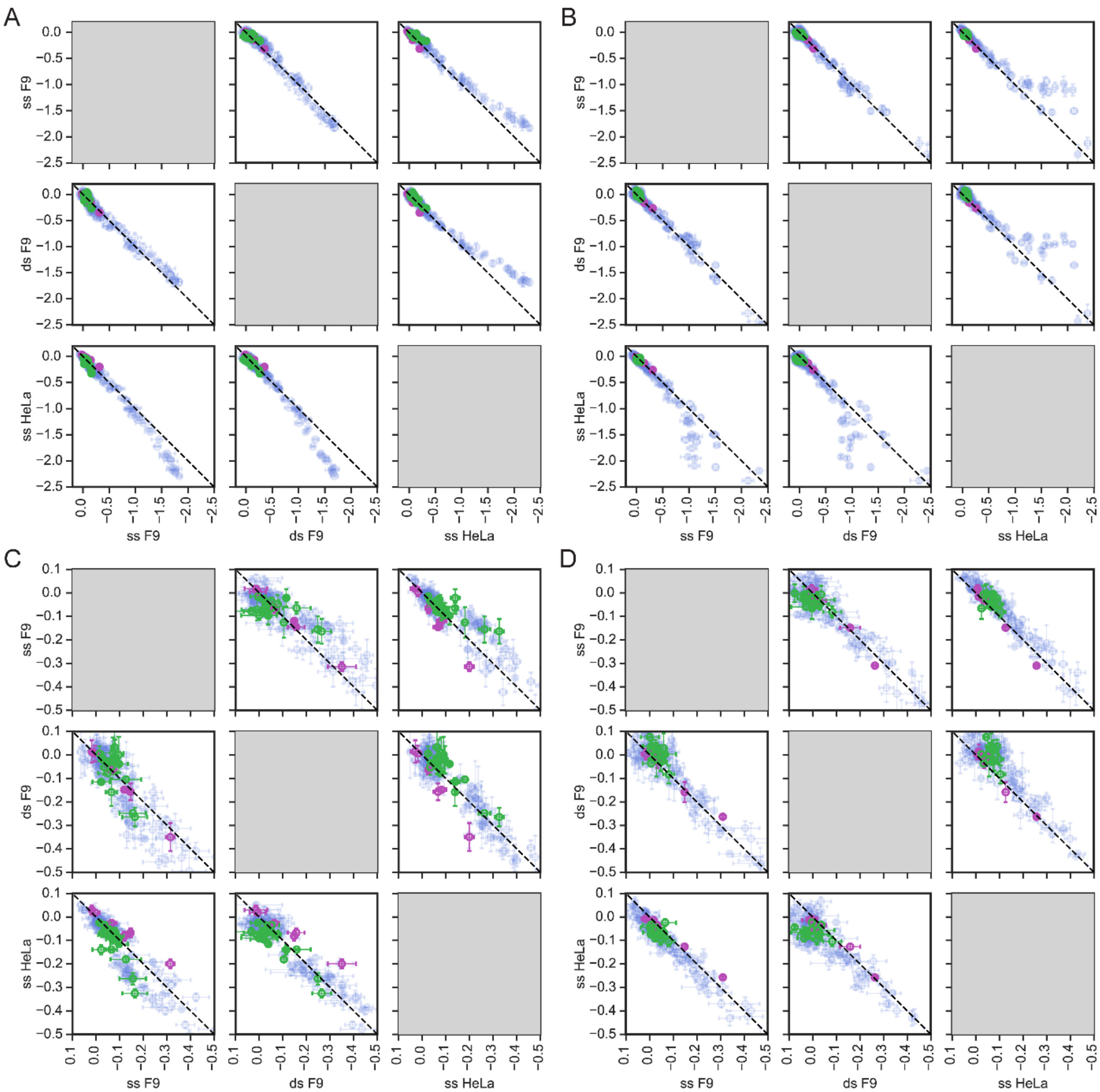


**Figure S7. Repression efficacies of equivalent sites were similar across biological replicates and cell types.** (A) Correlation of repression efficacies of miR-155 sites across biological replicates and cell lines. Plotted are pairwise comparisons of repression of reporter mRNAs containing miR-155 target sites (purple, canonical sites; blue, compound sites) in response to miR-155 transfection. Labels are as follows: ss F9, single-site repression efficacies from single-site MPRA in F9 cells; ds F9, single-site repression efficacies from MPRA that also examined dual sites in F9 cells; ss HeLa, single-site repression efficacies from single-site MPRA in HeLa cells. For each site, steady-state repression in each experiment is reported as mean ± standard error of the mean for the $N$ = 2 replicates from that experiment. (B) Correlation of repression efficacies of miR-124 sites across biological replicates and cell lines. Otherwise this panel is as in (A). (C) Zoomed-in view of (A). (D) Zoomed-in view of (B).

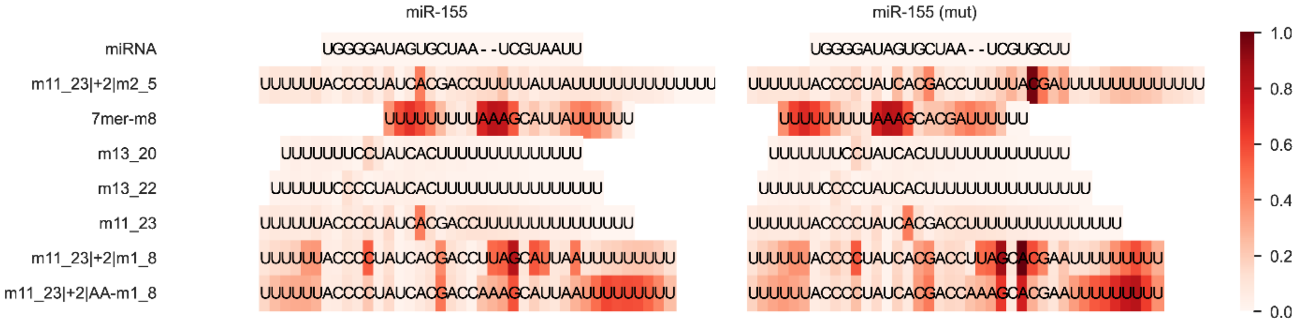


**Figure S8. Predicted structuredness of target RNAs used in experiments that measured association rates.** Folding of target RNAs was predicted using RNAfold with partition function, and the summed probability of being involved in a base pair was calculated from the ensemble for each position in the target RNA (8). Shading indicates the summed value calculated for each position, as indicated by the key.

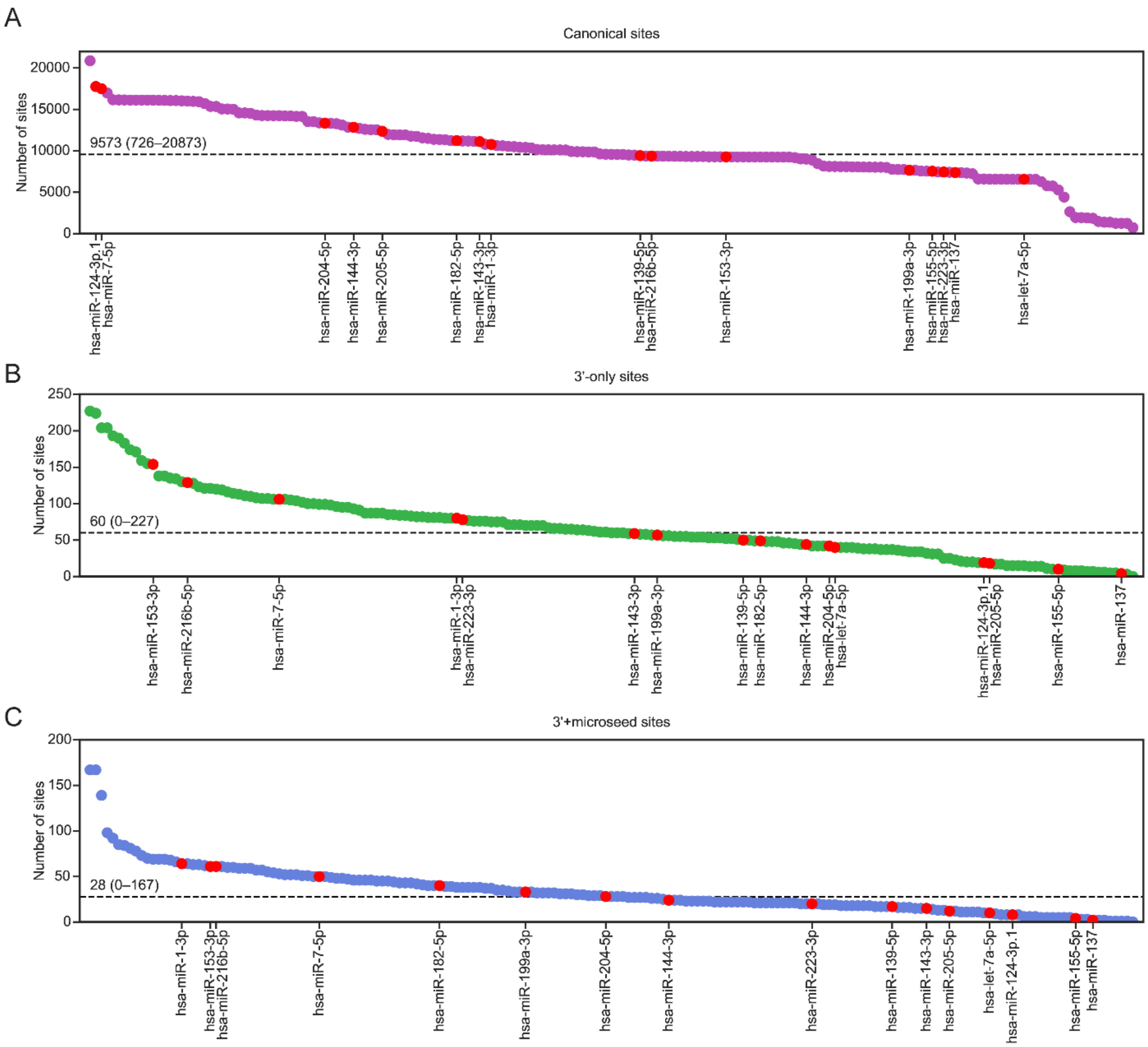


**Figure S9. Abundance of miRNA target sites in endogenous 3′ UTRs.** (A) Abundance of canonical sites in 3′ UTRs. Canonical sites were defined as 6mer, 7mer-A1, 7mer-m8, and 8mer sites, with or without additional 3′ complementarity. Plotted is the number of canonical sites found in endogenous 3′ UTRs for each of the 183 human miRNAs from the 108 seed families conserved to zebrafish. Note that for seed families with more than one member in humans, the same results are shown for each of the members. Red points highlight the results for the 16 miRNAs for which mRNA-seq data have been acquired after miRNA transfection, which are also labeled on the figure and analyzed in Figure S11. The black dashed line indicates the median; the values for the median and range are reported on the plot. (B) Abundance of 3′-only sites in 3′ UTRs. 3′-only sites were defined as those with ≥10 nucleotides of contiguous 3′ complementarity and <3 nucleotides of contiguous seed complementarity within a 20-nt window downstream of the 3' complementarity. Otherwise, this panel is as in panel (A). (C) Abundance of 3′+microseed sites in 3′ UTRs. 3′+microseed sites were defined as those with ≥10 nucleotides of contiguous 3′ complementarity and 3–5 nucleotides of contiguous seed complementarity within a 20-nt window downstream of the 3' complementarity. Otherwise, this panel is as in panel (A).

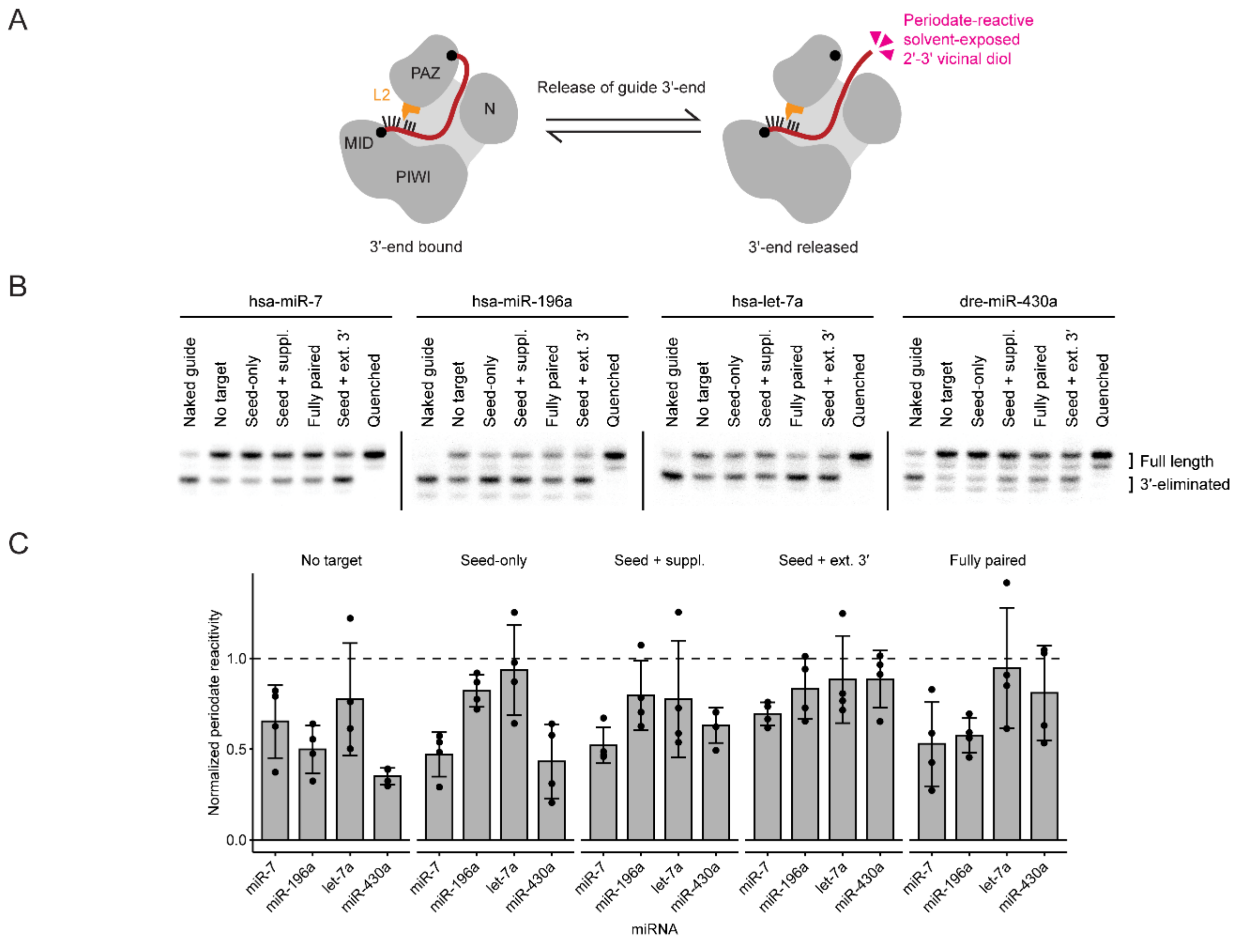


**Figure S10. 3′-end release from PAZ occurs even in the absence of target.** (A) Schematic of periodate probing assay to measure dissociation of the 3′-end from its binding pocket in the PAZ domain of AGO–miRNA complexes. (B) Representative gel images depicting the relative abundance of 3′-eliminated and full-length guide in AGO–miRNA complexes that contain either hsa-miR-7, hsa-miR-196a, hsa-let-7a, or dre-miR-430a after periodate probing. AGO-miRs were pre-saturated with either buffer-only negative control (no target), seed-only canonical target (seed-only), 3′-supplementary target (seed + suppl.), target with extensive 3′ complementarity and full seed complementarity (Seed + ext. 3′), or fully complementary target (fully paired). Included also is a pre-quenched negative control in which the periodate probing mixture was quenched before it was added to the binding reaction. (C) Quantification of relative abundance of 3′-eliminated product from gel images as in (B) for three replicates.

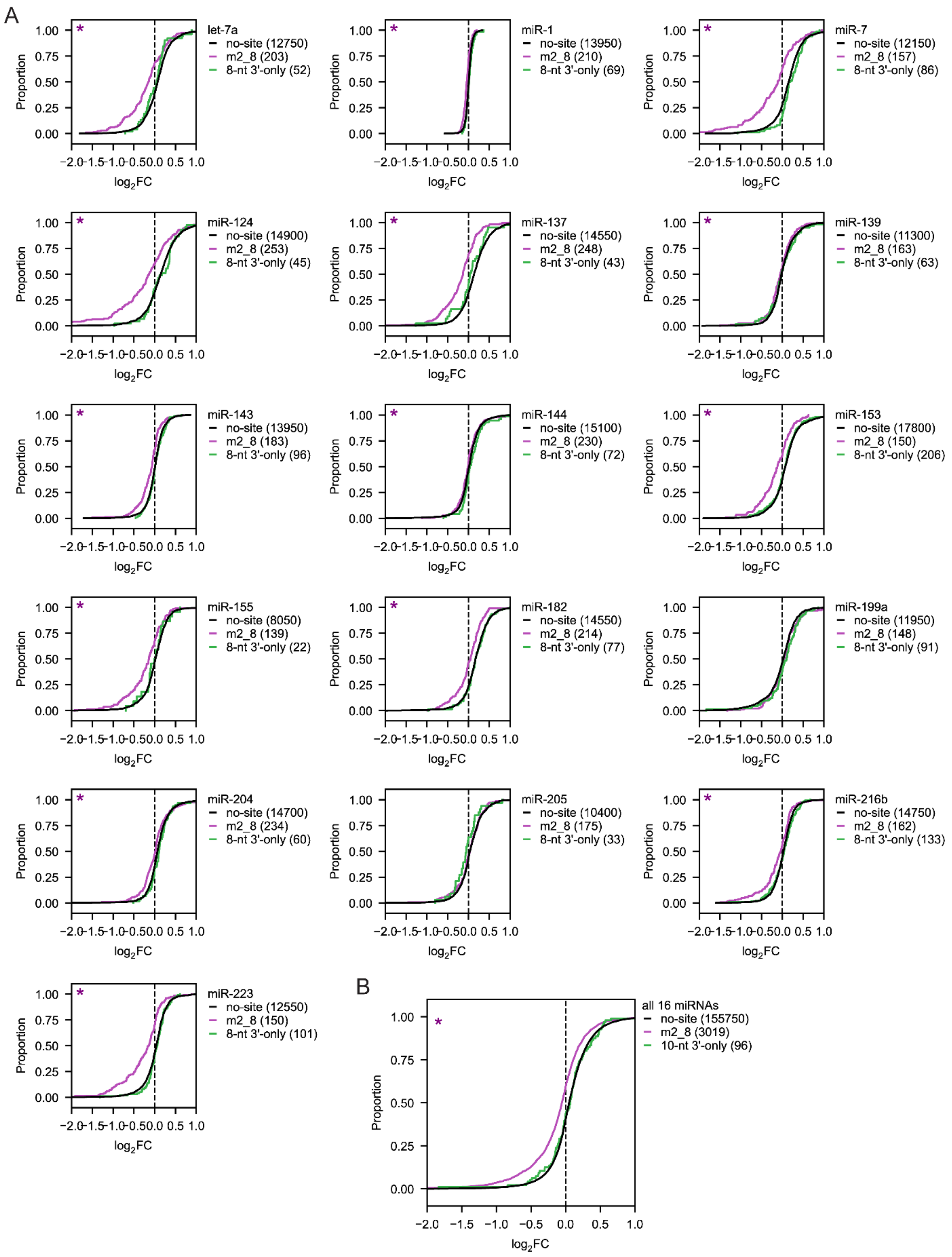
A
let-7a
no-site (12750)
m2_8 (203)
8-nt 3'-only (52)
miR-1
no-site (13950)
m2_8 (210)
8-nt 3'-only (69)
miR-7
no-site (12150)
m2_8 (157)
8-nt 3'-only (86)
miR-124
no-site (14900)
m2_8 (253)
8-nt 3'-only (45)
miR-137
no-site (14550)
m2_8 (248)
8-nt 3'-only (43)
miR-139
no-site (11300)
m2_8 (163)
8-nt 3'-only (63)
miR-143
no-site (13950)
m2_8 (183)
8-nt 3'-only (96)
miR-144
no-site (15100)
m2_8 (230)
8-nt 3'-only (72)
miR-153
no-site (17800)
m2_8 (150)
8-nt 3'-only (206)
miR-155
no-site (8050)
m2_8 (139)
8-nt 3'-only (22)
miR-182
no-site (14550)
m2_8 (214)
8-nt 3'-only (77)
miR-199a
no-site (11950)
m2_8 (148)
8-nt 3'-only (91)
miR-204
no-site (14700)
m2_8 (234)
8-nt 3'-only (60)
miR-205
no-site (10400)
m2_8 (175)
8-nt 3'-only (33)
miR-216b
no-site (14750)
m2_8 (162)
8-nt 3'-only (133)
miR-223
no-site (12550)
m2_8 (150)
8-nt 3'-only (101)
Proportion
log2FC
B
all 16 miRNAs
no-site (155750)
m2_8 (3019)
10-nt 3'-only (96)
Proportion
log2FC

**Figure S11. Attempts to detect repression of endogenous mRNAs containing single 3′-only sites were unsuccessful.** (A) Response of endogenous mRNAs containing either no site, a canonical 7mer-m8 site, or a ≥8-nt 3′-only site to a transfected miRNA (1). Cumulative distributions plot $\log_2$(fold-changes) after miRNA transfection for endogenous mRNAs containing either a single m2_8 (7mer-m8) canonical site (purple) or a single ≥8-nt 3′-only site (green) to the transfected miRNA in their 3′ UTR, comparing to control mRNAs with similar 3′ UTR length but no sites to the transfected miRNA. Purple asterisk (top-left corner) indicates that the distribution of fold changes for mRNAs containing m2_8 canonical sites was statistically different from that for the mRNAs containing no site (two-sided Kolmogorov–Smirnov test, $p < 0.05$). Green asterisk (top-left corner) indicates the same finding for the distribution of fold changes for mRNAs containing 3′-only sites. (B) Response of endogenous mRNAs containing either no site, a canonical 7mer-m8 site, or a ≥10-nt 3′-only site to a transfected miRNA, aggregated across 16 miRNA transfections. Cumulative distribution plots are as in (A) but aggregated across 16 miRNA transfections.

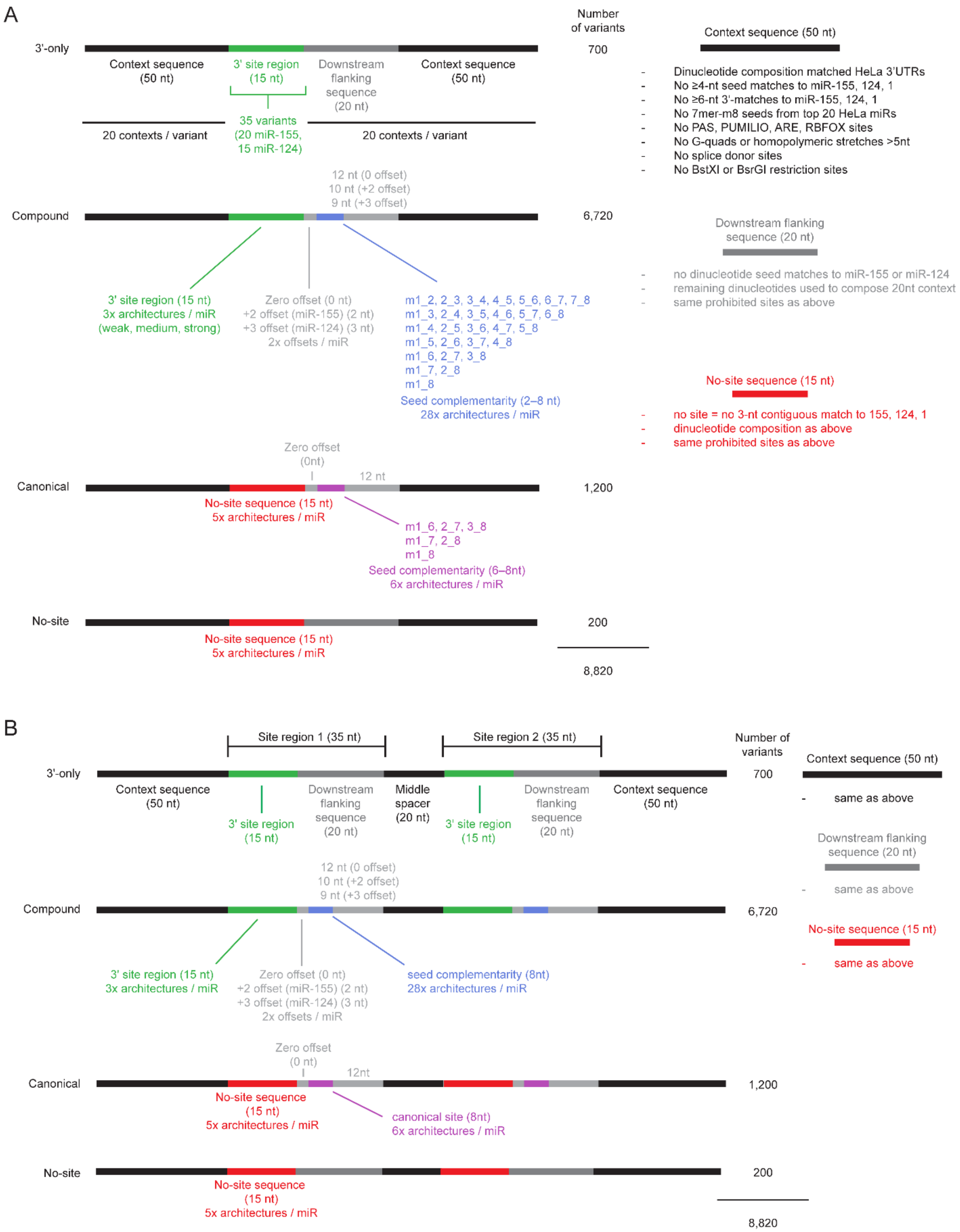
A
Number of variants
3'-only
Context sequence (50 nt)
3' site region (15 nt)
Downstream flanking sequence (20 nt)
Context sequence (50 nt)
700
35 variants (20 miR-155, 15 miR-124)
20 contexts / variant
20 contexts / variant
Context sequence (50 nt)
- Dinucleotide composition matched HeLa 3'UTRs
- No ≥4-nt seed matches to miR-155, 124, 1
- No ≥6-nt 3'-matches to miR-155, 124, 1
- No 7mer-m8 seeds from top 20 HeLa miRs
- No PAS, PUMILIO, ARE, RBFOX sites
- No G-quads or homopolymeric stretches >5nt
- No splice donor sites
- No BstXI or BsrGI restriction sites
12 nt (0 offset)
10 nt (+2 offset)
9 nt (+3 offset)
Compound
6,720
Downstream flanking sequence (20 nt)
- no dinucleotide seed matches to miR-155 or miR-124
- remaining dinucleotides used to compose 20nt context
- same prohibited sites as above
3' site region (15 nt)
3x architectures / miR
(weak, medium, strong)
Zero offset (0 nt)
+2 offset (miR-155) (2 nt)
+3 offset (miR-124) (3 nt)
2x offsets / miR
m1_2, 2_3, 3_4, 4_5, 5_6, 6_7, 7_8
m1_3, 2_4, 3_5, 4_6, 5_7, 6_8
m1_4, 2_5, 3_6, 4_7, 5_8
m1_5, 2_6, 3_7, 4_8
m1_6, 2_7, 3_8
m1_7, 2_8
m1_8
Seed complementarity (2–8 nt)
28x architectures / miR
No-site sequence (15 nt)
- no site = no 3-nt contiguous match to 155, 124, 1
- dinucleotide composition as above
- same prohibited sites as above
Zero offset (0nt)
12 nt
Canonical
1,200
No-site sequence (15 nt)
5x architectures / miR
m1_6, 2_7, 3_8
m1_7, 2_8
m1_8
Seed complementarity (6–8nt)
6x architectures / miR
No-site
200
No-site sequence (15 nt)
5x architectures / miR
8,820
B
Site region 1 (35 nt)
Site region 2 (35 nt)
Number of variants
3'-only
700
Context sequence (50 nt)
Downstream flanking sequence (20 nt)
Middle spacer (20 nt)
3' site region (15 nt)
Downstream flanking sequence (20 nt)
Context sequence (50 nt)
3' site region (15 nt)
Context sequence (50 nt)
- same as above
Downstream flanking sequence (20 nt)
- same as above
12 nt (0 offset)
10 nt (+2 offset)
9 nt (+3 offset)
Compound
6,720
No-site sequence (15 nt)
- same as above
3' site region (15 nt)
3x architectures / miR
Zero offset (0 nt)
+2 offset (miR-155) (2 nt)
+3 offset (miR-124) (3 nt)
2x offsets / miR
seed complementarity (8nt)
28x architectures / miR
Zero offset (0 nt)
12nt
Canonical
1,200
No-site sequence (15 nt)
5x architectures / miR
canonical site (8nt)
6x architectures / miR
No-site
200
No-site sequence (15 nt)
5x architectures / miR
8,820

**Figure S12. Detailed schematic of library designs for MPRAs.** (A) Design of single-site MPRA library inserts. (B) Design of dual-site MPRA library inserts.

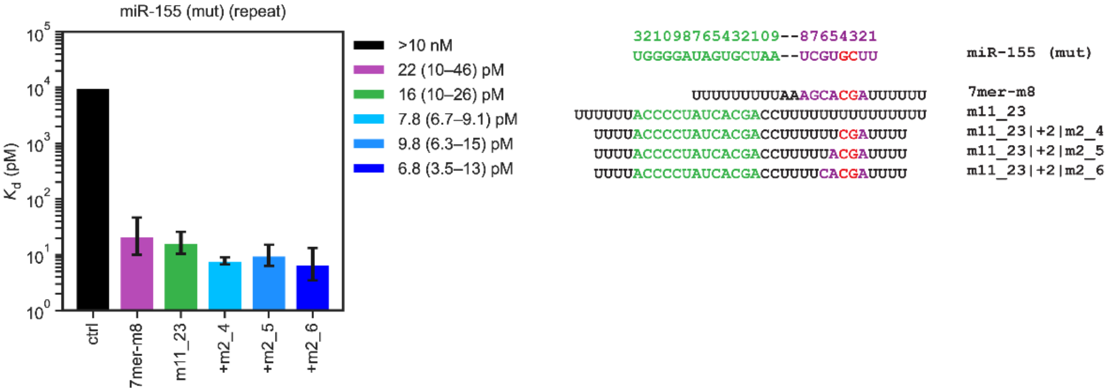


**Figure S13. Repeat of equilibrium binding assays of Figure 2C with a shorter target RNA.** Equilibrium binding assays were as in Figure 2C for miR-155 (mut) targets containing m11_23 3′ complementarity in combination with different segments of 2–5-nt seed complementarity at +2 offset. Target with m11_23|+2|m2_5 pairing in Figure 2C had slightly longer poly(U) flanking sequence than other targets, whereas the m11_23|+2|m2_5 target used here was equal in length to other targets, and bound about as well as m11_23|+2|m2_4 and m11_23|+2|m2_6, as expected.
